# Sex-specific T cell exhaustion drives differential immune responses in glioblastoma

**DOI:** 10.1101/2022.08.17.503211

**Authors:** Juyeun Lee, Michael Nicosia, Daniel J. Silver, Cathy Li, Defne Bayik, Dionysios C. Watson, Adam Lauko, Sadie Johnson, Mary McGraw, Matthew M. Grabowski, Danielle D. Kish, Amar Desai, Wendy Goodman, Scott J. Cameron, Hideo Okada, Anna Valujskikh, Robert L. Fairchild, Manmeet S. Ahluwalia, Justin D. Lathia

**Affiliations:** Department of Cardiovascular and Metabolic Sciences, Lerner Research Institute, Cleveland Clinic, Cleveland, OH; Department of Infection and Immunity, Lerner Research Institute, Cleveland Clinic, Cleveland, OH; Case Comprehensive Cancer Center, Cleveland, OH; Hematology/Oncology Division, Department of Medicine, University Hospitals Cleveland Medical Center; Medical Scientist Training Program, Department of Medicine, Case Western Reserve University, Cleveland OH; Rose Ella Burkhardt Brain Tumor Center, Cleveland Clinic, Cleveland, OH; Department of Medicine, Case Western Reserve University, Cleveland OH; Department of Neurological Surgery, University of California San Francisco, San Francisco, CA; Miami Cancer Institute, Miami, FL

## Abstract

Sex differences in glioblastoma (GBM) incidence and outcome are well recognized, and emerging evidence suggests that these extend to genetic/epigenetic and cellular differences, including immune responses. However, the mechanisms driving immunological sex differences are not fully understood. Using GBM models, we demonstrate that T cells play a critical role in driving GBM sex differences. Male mice exhibited accelerated tumor growth, with decreased T cell infiltration and increased T cell exhaustion. Furthermore, a higher frequency of progenitor exhausted T cells was found in males, with improved responsiveness to anti-PD1 treatment. Bone marrow chimera and adoptive transfer models indicated that T cell-mediated tumor control was predominantly regulated in a cell-intrinsic manner, which was further corroborated by *in vitro* exhaustion assays. Moreover, increased T cell exhaustion was observed in male GBM patients. These findings demonstrate sex-specific pre-determined behavior of T cells is critical in inducing sex differences in GBM progression and immunotherapy response.

**Statement of significance:** Immunotherapies in GBM patients have been unsuccessful due to a variety of factors including the highly immunosuppressive tumor microenvironment in GBM. This study demonstrates that sex-specific T cell behaviors are predominantly intrinsically regulated, further suggesting sex-specific approaches can be leveraged to potentially improve therapeutic efficacy of immunotherapy in GBM.

## Introduction

Glioblastoma (GBM) is the most common primary malignant brain tumor, and patients with GBM experience poor prognosis despite aggressive current standard-of-care therapies including surgical resection, radiotherapy, and chemotherapy with temozolomide (1). One reason that GBM is difficult to treat is its highly immunosuppressive tumor microenvironment (TME). GBM tumors are infiltrated with suppressive myeloid populations including macrophages, myeloid-derived suppressor cells (MDSCs), and microglia (2). A reduction in T cells in the circulation further contributes to poor anti-GBM immunity, with sequestration of naïve T cells in bone marrow and involution of primary and secondary lymphoid organs suggested as mechanisms of T cell reduction (3, 4). Clinical trials in GBM employing immune checkpoint inhibitors (ICIs) such as anti-PD1 and anti-CTLA4 monoclonal antibodies have generally not shown significant improvement in overall survival, even when used in combination therapies with existing treatment options (5-7), with an exception of a single trial that demonstrated a modest but statistically significant survival benefit when ICI was employed in the neoadjuvant setting (8). Given the unique TME and anatomical immune privilege in GBM, there is a pressing need to understand how to reinvigorate immune responses in GBM.

Adding to the complexity of GBM treatment is a sex difference in disease outcome, with male patients exhibiting a 1.6-fold higher incidence and poorer prognosis after treatment compared to female patients (9). While tumor-intrinsic factors underlying sex differences have been identified in GBM (10, 11), sex differences in anti-tumor immunity may also contribute. In general, females exhibit stronger immune responses than males, as mostly shown in autoimmune and infectious diseases, and the differences are attributed to sex hormones and/or sex chromosomes (12). Overall, male-biased prevalence of cancers in non-reproductive organs have been reported (13), yet the underlying mechanisms remain to be elucidated.

In many tumors, T cell function is disrupted, and addressing this has been the focus of many immunotherapies. T cell exhaustion refers to a dysfunctional state of T cells that is characterized by high expression of inhibitory receptors, poor effector function, and decreased proliferative potential and is mediated by epigenetic remodeling (14). Chronic antigen stimulation in infection and cancers induces T cell exhaustion, resulting in impaired control of disease. Exhausted T cells are comprised of two distinct subsets - stem-like/progenitor exhausted T cells with self-renewal properties, which then transition to the irreversible stage of terminally exhausted T cells. Progenitor exhausted T cells can temporarily differentiate into “effector-like” T cells in response to anti-PD1 blockade that effectively inhibit tumor growth (15, 16). Male-biased T cell exhaustion was observed in a variety of cancers including melanoma and bladder cancer (17, 18), demonstrating that sex differences in T cell exhaustion lead to divergent disease outcomes in males and females. Whilst GBM is highly enriched with exhausted T cells (19), it remains unknown the extent to which sex-specific T cell exhaustion mediates the sex differences in GBM survival.

We previously demonstrated sex-specific behaviors of MDSC subsets in GBM (20). In this study, we hypothesized that the sex differences in response to GBM extend beyond the myeloid lineage and here, report sex differences in T cell exhaustion that underlie differential ICI response in GBM.

## Results

### T cells are a critical driver of sex differences in GBM

We have previously shown sex differences in survival using the syngeneic mouse glioma models SB28 and GL261 (20), with male mice experiencing a worse outcome than female mice. To investigate the role of immune cell populations in this sex-based survival difference, we utilized immunocompromised mouse strains with different degrees of immunodeficiency. Intracranial tumor implantation revealed that the sex-specific survival difference observed in wild-type mice was not present in immune-deficient NSG mice (**Fig. 1A**). For these survival assessments, we used a reduced number of tumor cells in NSG mice (SB28-10,000 cells/mouse, GL261-20,000 cells/mouse) compared to wild-type mice (SB28-15,000 cells/mouse, GL261-25,000 cells/mouse) due to accelerated tumor growth in immune-deficient models. We also did not observe any difference in survival of NSG mice challenged with a lower number of tumor cells (**Supplementary Fig. 1A**). To further specify the immune cell population responsible for the survival difference observed in immunocompetent B6 mice, we used RAG1^-/-^ mice that lack only mature T and B lymphocytes, while other innate immune cells remain intact. We observed no sex difference in survival outcomes in tumor-bearing RAG1^-/-^ mice (**Fig. 1A**), suggesting a role for lymphocytes. The male-biased aggressive tumor growth was not due to elevated female immune responses against male-specific antigens expressed by the tumor cells (SB28 and GL261), as we confirmed that neither tumor cell line contains a Y chromosome (21) or expresses Y chromosome-encoded genes, particularly H-Y minor histocompatibility antigen encoding gene (*smcy*) (**Supplementary Fig. 1B-C**).

**Figure 1.**
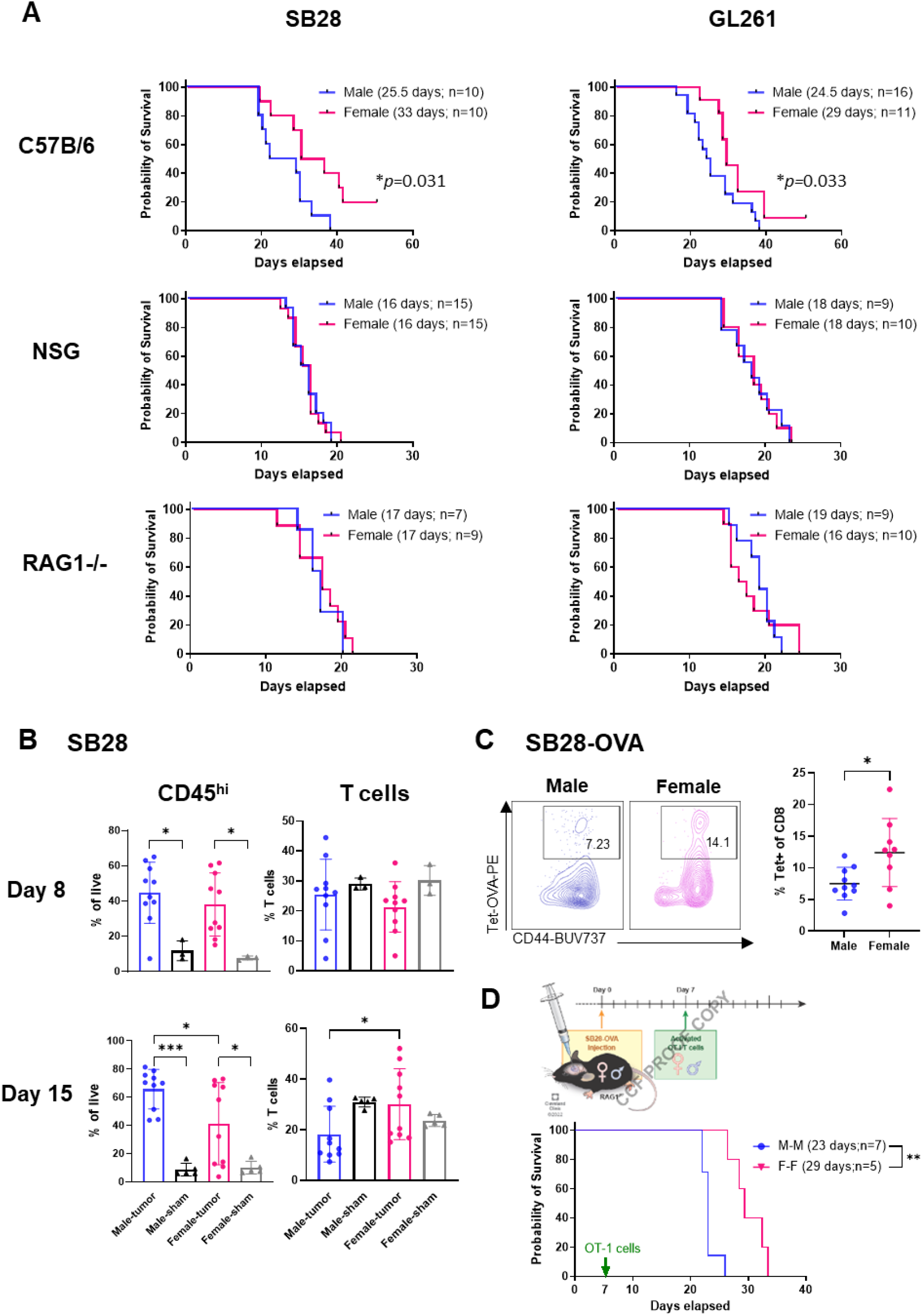
T cells drive sex differences in GBM survival. (A) Survival analysis was performed after intracranial injection of mouse GBM cell lines SB28 and GL261 in immunocompetent B6 mice (SB28-15,000 cells/mouse, GL261-25,000 cells/mouse) and immune-deficient NSG (SB28-10,000 cells/mouse, GL261-20,000 cells/mouse) and RAG1^-/-^ mice (SB28-15,000 cells/mouse, GL261-25,000 cells/mouse). Median survival days and number of animals are indicated in the graph legend. Data combined from two to three independent experiments. Statistical significance was determined by log-rank test, considering *p*-value <0.05 to be significant. (B) Frequency of CD45^hi^ immune cells and CD3^+^ T cells from the tumor-bearing left hemisphere of SB28-injected mice or the left hemisphere of the sham-injected group on day 8 and 15. Data shown as mean±SD from two independent experiments. N=10 for SB28-bearing mice and n=3-5 for sham-injected mice. One-way ANOVA with Tukey’s multiple comparison test was performed to determine statistical significance (\**p*<0.05, \*\**p*<0.01, \*\*\**p*<0.001). (C) Frequency of OVA-specific CD8^+^ T cells were measured using tetramer antibody from the tumor-bearing left hemisphere of SB28-OVA (25,000 cells/mouse)-injected B6 wild-type mice on day 14 post-tumor implantation. Data shown as mean±SD from two independent experiments. N=9/group. \**p*<0.05 was determined by unpaired unpaired *t*-test. (D) Survival analysis was performed after adoptive transfer of OT-I cells into RAG1^-/-^ mice bearing SB28-OVA tumors. Median survival days and number of animals are indicated in the graph legend. Data combined from two independent experiments. Log-rank test was performed and \*\**p*<0.01 was determined to be significant.

To understand immunological differences between male and female hosts, we profiled tumor-infiltrating immune cells using flow cytometry at two different time points (**Fig. 1B, Supplementary Fig. 2A**). On day 8 post-tumor implantation, when T cells are expected to be fully primed and activated, there was increased CD45^hi^ immune cell infiltration with tumor compared to the sham group. Males and females showed comparable levels of total immune cells as well as T cells at this early time point. On day 15 post-tumor implantation, when some mice start to show neurological symptoms as a result of an advanced tumor, we found that CD45^hi^ immune cells were significantly higher in males compared to females. Further analysis revealed that immune cells in female tumors were more enriched in T cells (**Fig. 1B**), whereas male tumors had a higher ratio of macrophages (**Supplementary Fig. 2B**). We did not observe any major differences in other immune cell subsets on either day 8 or day 15 **(Supplementary Fig. 2B**).

To test whether sex differences in the diversity of the T cell receptor (TCR) repertoire affect T cell responsiveness to tumor antigens, we evaluated antigen-specific T cell responses by measuring the ovalbumin (OVA)-specific CD8^+^ T cell population using a tetramer antibody after implantation of OVA-expressing tumor cells (SB28-OVA) into male and female wild-type B6 mice. Similar to polyclonal T cells (**Fig. 1B**), OVA-specific CD8^+^ T cells were significantly higher in female mice (**Fig. 1C**). These data indicate that female T cells have the potential to be more reactive to the tumor antigen regardless of TCR diversity.

To further investigate the extent to which the sex difference in survival was driven by T cells, we transferred activated OT-I T cells into SB28-OVA tumor-bearing RAG1^-/-^ mice. T cells were indeed responsible, in part, for the sex difference in survival, as male mice receiving male T cells exhibited significantly faster tumor growth compared to female mice receiving female T cells (**Fig. 1D**), recapitulating the survival differences observed in the immunocompetent model (**Fig. 1A**). Taken together, these data suggest that T cells play an essential role in the sex differences in survival observed in mouse GBM models.

### Male T cells become exhausted more quickly than female T cells, with a higher frequency of progenitor exhausted T cell subsets

We next performed extensive immune cell profiling focused on T cells to understand how T cells underlie the sex differences in GBM progression. Increased CD8^+^ and CD4^+^ T cell populations were observed in female tumors compared to male tumors on day 15 post-tumor implantation, while the frequency of Foxp3^+^ cells was comparable, indicating that the differences are likely from effector T cells (**Fig. 2A**). Next, we asked whether female T cells were primed and activated earlier, thereby more effectively attenuating tumor growth. To address this possibility, we evaluated activated T cell phenotypes in the draining lymph nodes at earlier time points after tumor implantation (**Supplementary Fig. 3A**). No differences in phenotype or functionality were observed between male and female T cells in draining lymph nodes, indicating that differential kinetics of T cell activation was likely not the reason for the sex differences in T cell infiltration into tumors.

**Figure 2.**
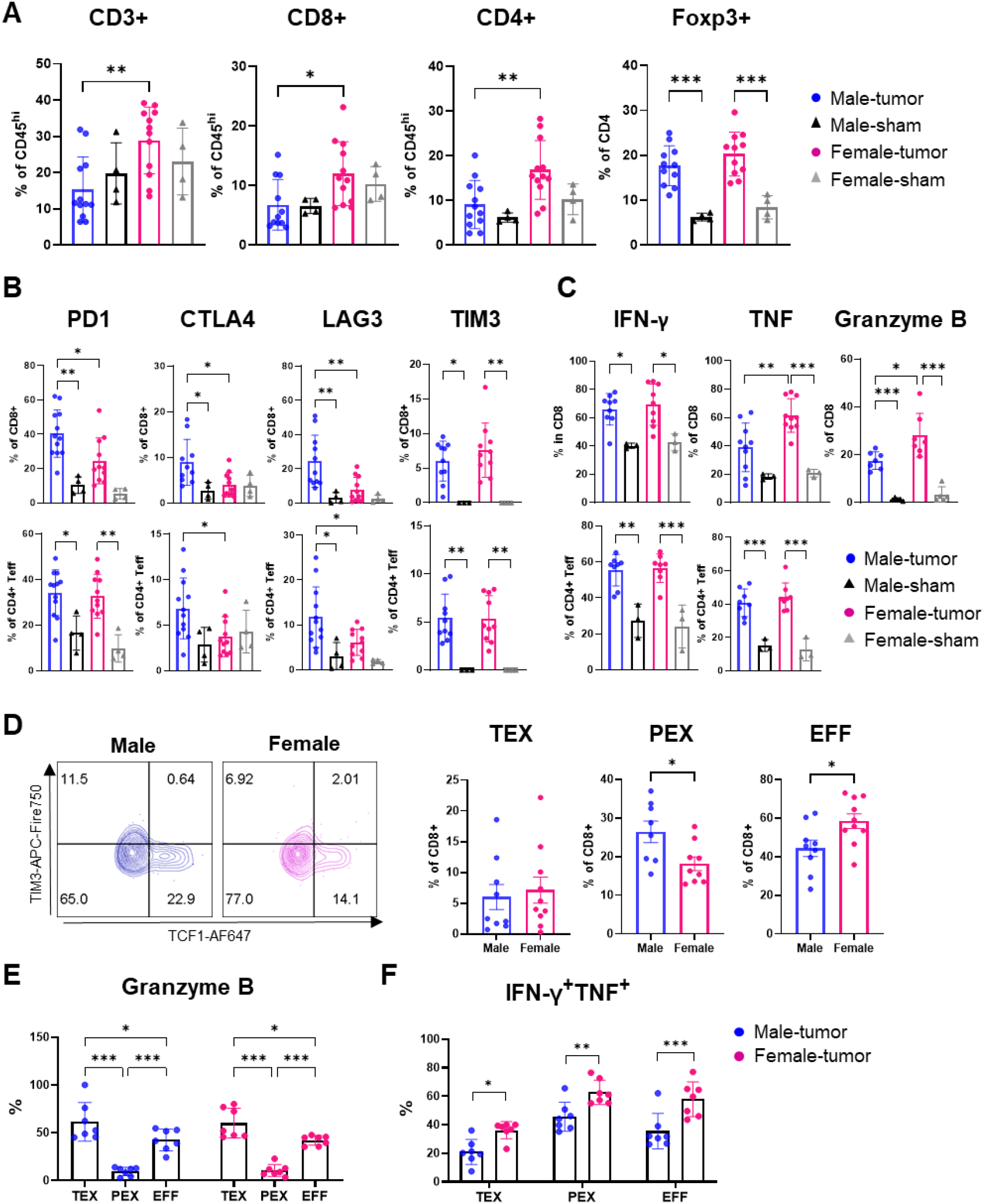
More male CD8^+^ T cells are functionally exhausted and skewed toward a progenitor exhausted T cell phenotype. Tumor-infiltrating T cells were analyzed on day 15 post-implantation of SB28 tumor cells. (A) Frequency of T cell subsets in CD45^hi^ immune cells. Data combined from two independent experiments. n=11-12 for the SB28-injected group and n=4 for the sham-injected group. (B) Inhibitory receptor expression in CD8^+^ and CD4^+^Foxp3^-^ effector T cells (Teff). Data combined from two independent experiments. n=10-12 for the SB28-injected group and n=4 for the sham-injected group. (C) Intracellular cytokine expression in CD8^+^ and CD4^+^Foxp3^-^ effector T cells was measured after ex vivo stimulation. Data combined from two independent experiments. n=7-10 for the SB28-injected group and n=3 for the sham-injected group. (D) Exhausted T cell subsets in CD8^+^ T cells: TEX (terminally exhausted; CD8^+^CD44^+^PD1^+^TCF1^-^TIM3^+^), PEX (progenitor exhausted; CD8^+^CD44^+^PD1^+^TCF1^+^TIM3^-^), EFF (effector; CD8^+^CD44^+^TCF1^-^TIM3^-^). Data combined from two independent experiments. n=9-10 for the SB28-injected group and n=4 for the sham-injected group. Intracellular expression of (E) granzyme B and (F) IFN-γ^+^TNF^+^ in each CD8^+^ T cell subset after *ex vivo* stimulation. Data combined from two independent experiments. n=11-12 for SB28-injected group and n=4 for sham-injected group. Two-way ANOVA analysis with Tukey’s multiple comparison test (A-C, E,F) or unpaired Student’s *t* test (D) was performed (\**p*<0.05, \*\**p*<0.01, \*\*\**p*<0.001).

Further analysis revealed that male CD8^+^ T cells express a higher frequency of inhibitory receptors, including PD1, CTLA4, and LAG3 but not TIM3, compared to female CD8^+^ T cells (**Fig. 2B**). Additionally, male CD8^+^ T cells showed decreased levels of intracellular cytokine expression compared to female CD8^+^ T cells following *ex vivo* stimulation (**Fig. 2C**). CD4^+^Foxp3^-^ effector T cells showed differences such as expression of CTLA4 and LAG3, but these differences were not as prominent as in CD8^+^ T cells. The differences were not observed at the early time point (day 8, **Supplementary Fig. 3B-D**), consistent with the findings in **Fig. 1B**. We also confirmed these findings using another GBM model, GL261, which showed no clear difference in T cell frequency but differences in inhibitory receptor expression as well as TNF expression (**Supplementary Fig. 4A-C**). These phenotypic differences were only observed in T cells infiltrating tumors, as no significant changes in inhibitory receptor and cytokine expression were found in T cells from blood and bone marrow (**Supplementary Fig. 5**).

The increased expression of inhibitory receptors and decreased cytokine expression in male CD8^+^ T cells prompted us to hypothesize that male and female CD8^+^ T cells exhibit differential exhaustion status. It is well established that exhausted CD8^+^ T cells are comprised of two subsets, stem-like/progenitor exhausted CD8^+^T cells (**PEX**; CD8^+^CD44^+^PD1^+^TCF1^+^TIM3^-^) and terminally exhausted CD8^+^ T cells (**TEX**; CD8^+^CD44^+^PD1^+^TCF1^-^TIM3^+^) (16). Therefore, we evaluated the frequency of exhausted T cell subsets and effector T cells (**EFF**; CD8^+^CD44^+^TCF1^-^TIM3^-^) in tumor-infiltrating CD8^+^ T cells. Strikingly, male CD8^+^ T cells contained a significantly higher ratio of PEX compared to female CD8^+^ T cells, while female T cells had elevated frequencies of EFF (**Fig. 2D**). A higher frequency of the PEX subset in males was also observed in the GL261 model (**Supplementary Fig. 4D**). No difference was observed in the TEX population between male and female CD8^+^ T cells (**Fig. 2D**) as well as at the early time point (**Supplementary Fig. 3E**). Intracellular expression of granzyme B confirmed the gating strategy for exhausted T cell subsets, as TEX showed the highest level of granzyme B, while PEX showed minimal expression as described previously (16)(**Fig. 2E**). As exhausted CD8^+^ T cells have reduced capacity to produce multiple cytokines (16), we evaluated the proportion of cells simultaneously producing IFN-γ and TNF. Interestingly, all three subsets of female CD8^+^ T cells were more polyfunctional,, suggesting that the fundamental sex difference in T cell functionality exists independent of exhaustion status (**Fig. 2F**). Collectively, these data indicate that male and female T cells undergo exhaustion at different rates, with higher progenitor exhausted T cells in males whereas higher effector cytokine production in female CD8^+^ T cells, which may result in survival differences.

### Males benefit from anti-PD1 therapy more than females

Progenitor exhausted T cells are better able to control tumor growth and respond to anti-PD1 treatment (15, 16). Thus, we hypothesized that males will respond better than females to anti-PD1 monoclonal antibody (mAb) treatment due to the high frequency of progenitor cells in male tumors. To test this possibility, we treated male and female mice bearing SB28 tumors with anti-PD1 mAb or isotype antibody (**Fig. 3A**). In accordance with our prediction, anti-PD1 mAb treatment significantly extended the survival of male mice compared to the isotype-treated group (**Fig. 3B**). However, the treatment effect was mild in females, as previously reported (22).

**Figure 3.**
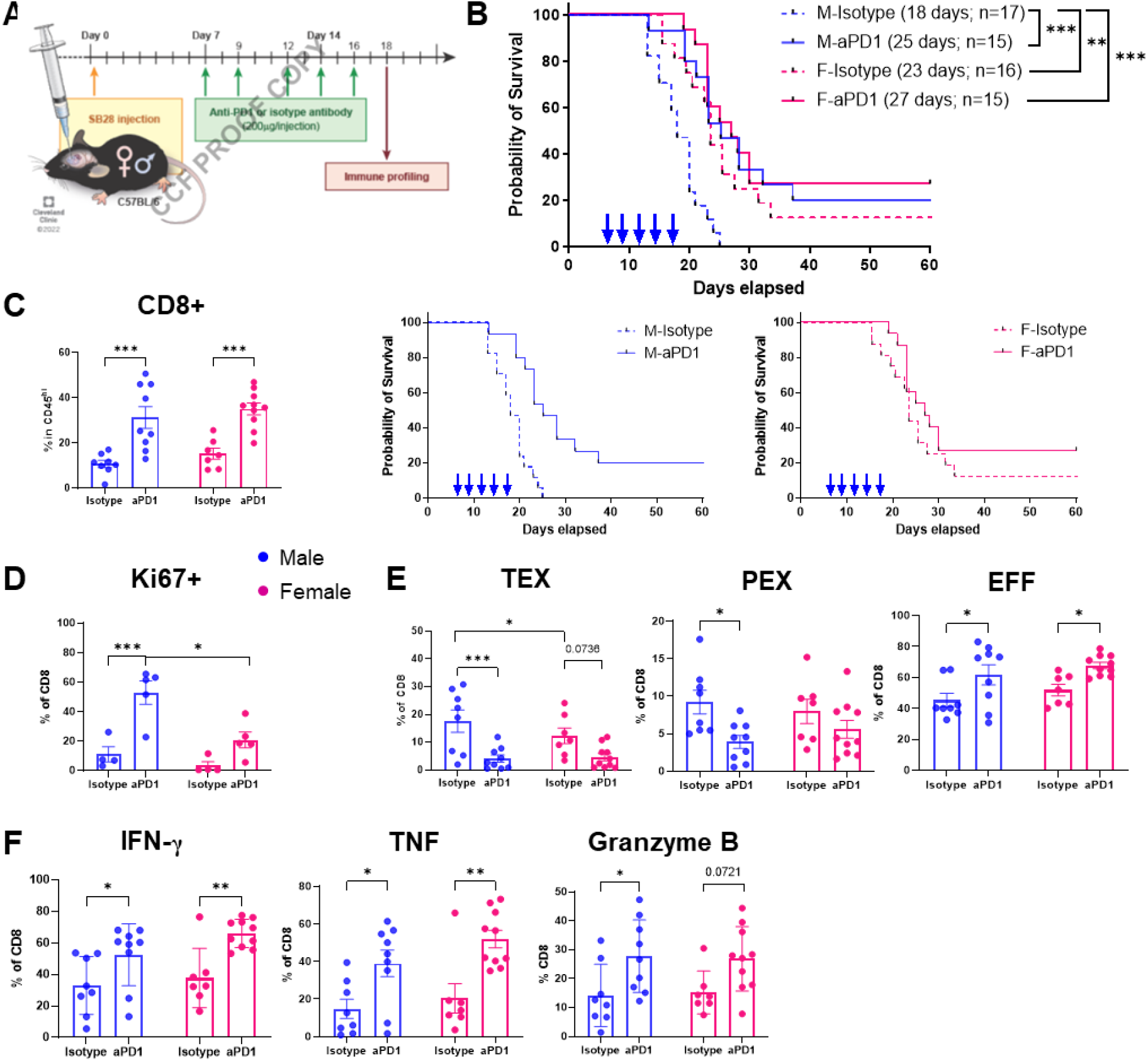
Males are more responsive to anti-PD1 therapy. (A) Schematics depicting treatment regimen for anti-PD1 and immune profiling. (B) Kaplan-Meier curves depicting survival of male and female mice treated with anti-PD1 or isotype antibodies (10 mg/kg) starting from day 7 post-intracranial tumor implantation. Combined results from three independent experiments with log-rank test (\*\**p*<0.01, \*\*\**p*<0.001). Median survival length and number of animals are indicated. (C-F) Immunophenotyping was performed on tumor-infiltrating immune cells on day 18 after the last treatment. Data combined from two independent experiments. n=9-10 for the anti-PD1-treatment group and n=7-8 for the isotype antibody-treated group. (C) Percentage of CD8^+^ T cells in CD45^hi^ cells. (D) Proliferation marker Ki-67 expression in CD8^+^ T cells. Data shown as mean±SD of n=5/group from one of two independently repeated experiments. (E) Frequency of exhausted T cell subsets in CD8^+^ T cells. (F) Percentages of intracellular CD8^+^ T cells expressing IFN-γ^+^, TNF^+^, and granzyme B. Two-way ANOVA analysis with Tukey’s multiple comparison test was performed (\**p*<0.05, \*\**p*<0.01, \*\*\**p*<0.001).

To further interrogate the survival differences between males and females in response to anti-PD1 mAb treatment, we analyzed tumor infiltrating T cells using flow cytometry two days after the last treatment. The frequency of tumor-infiltrating CD8^+^ T cells was increased in both males and females with anti-PD1 mAb treatment (**Fig. 3C**), while other immune cells in the tumor showed a similar trend regardless of sex (**Supplementary Fig. 6A**). PD1 expression was dramatically reduced in both CD8^+^ and CD4^+^ T cells with anti-PD1 mAb treatment, whereas CTLA4 and LAG3 expression was not altered (**Supplementary Fig. 6B**). Interestingly, we observed a significant increase of Ki67^+^ CD8^+^ T cells (**Fig. 3D**) and CD4^+^ effector T cells (**Supplementary Fig. 6C**) in males, suggesting that male T cells became more proliferative upon anti-PD1 mAb treatment. PD1 blockade also led to a significant decrease in the TEX and PEX subsets in males, whereas the decrease was moderate in females (**Fig. 3E**). Cytokine expression was elevated in both male and female CD8^+^ T cells (**Fig. 3F**), but not in CD4^+^ effector T cells (**Supplementary Fig. 6D)**, with anti-PD1 mAb treatment. Taken together, these results indicate that anti-PD1 blockade was more effective on male T cells with significant changes in reducing exhausted T cell subsets and increasing proliferation compared to females, which may lead to survival benefit upon treatment in males.

### Immune cell-intrinsic regulation of sex differences in GBM

Next, we asked whether the sex differences in T cell phenotypes and GBM survival were driven by a hematopoietic immune cell-intrinsic or cell-extrinsic manner. To test this, we generated sex-matched or mismatched bone marrow chimeras by transferring T cell-depleted bone marrow cells into lethally irradiated recipient mice (**Fig. 4A**). We depleted pre-existing T cells in the bone marrow to prevent an induction of graft-versus-host disease in female-to-male group as well as to newly generate T cells from the donor hematopoietic stem cells and fully matured in the recipients’ thymus **(Supplementary Fig. 7A**). The immune components of the recipients were successfully reconstituted by the donor cells after 6 weeks (CD45.1^+^ cells > 95%), and no significant difference in the frequency of circulating immune cells was observed at steady state (**Supplementary Fig. 7B**).

**Figure 4.**
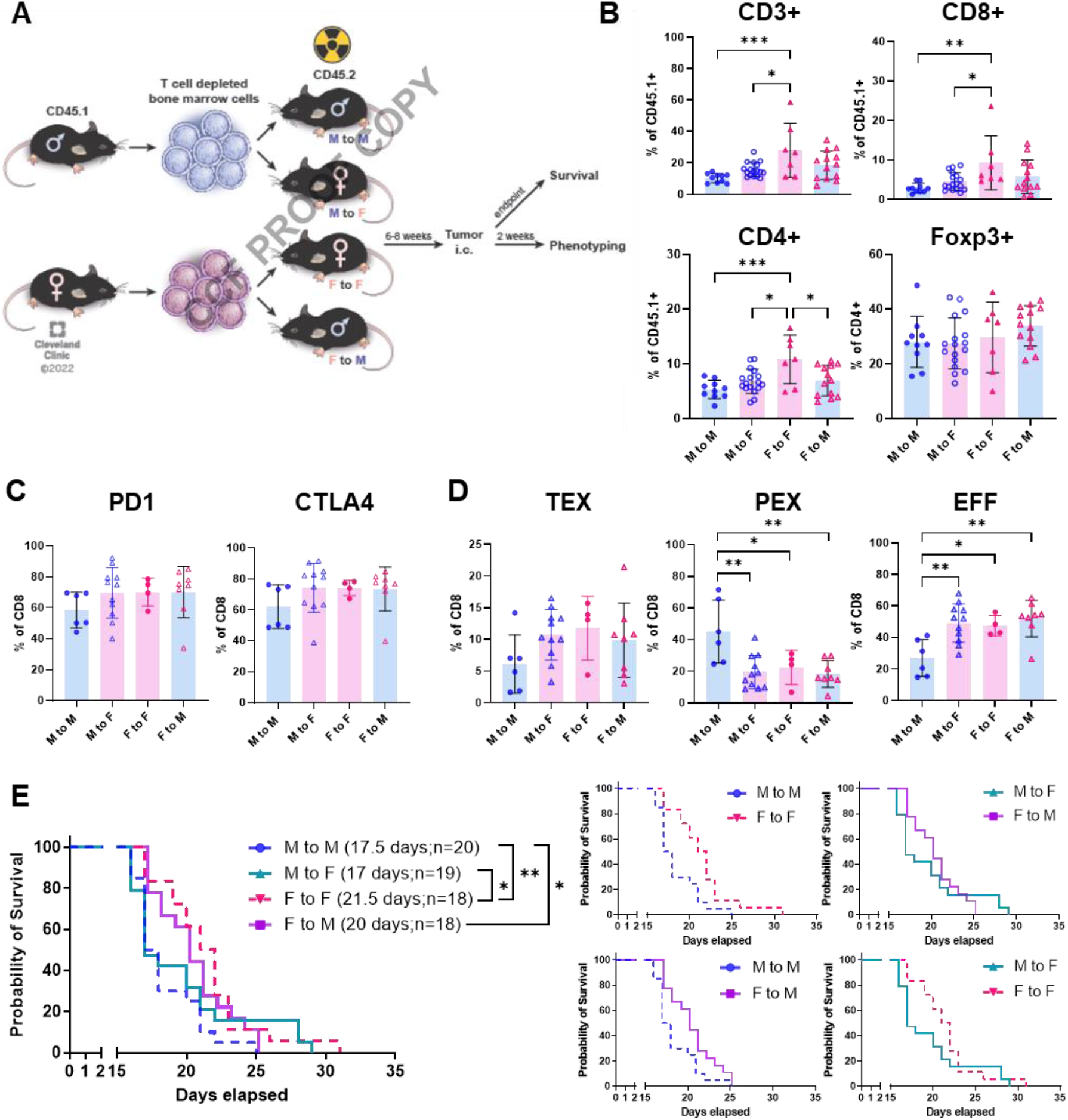
Immune cell-intrinsic and cell-extrinsic effect in GBM survival. (A) A schematic figure of the generation of bone marrow chimera models. Immune profiling was performed from the tumor-bearing hemisphere on day 14 post-tumor implantation (SB28, 10,000 cells/injection). (B) Frequency of T cell subsets in CD45^hi^ cells. Data combined from three independent experiments. n=7-17 per group. One-way ANOVA analysis with Tukey’s multiple comparison test (\**p*<0.05, \*\**p*<0.01, \*\*\**p*<0.001). (C) Percentage of PD1 and CTLA4 expression and (D) exhausted T cell subsets in CD8^+^ T cells. Data combined from three independent experiments. n=4-11 per group. One-way ANOVA analysis with Tukey’s multiple comparison test (\**p*<0.05, \*\**p*<0.01). (E) Kaplan-Meier curves depicting survival of bone marrow chimeras after intracranial injection of SB28 cells. Data shown is combined from two independent experiments. n=18=20 per group. Statistical significance was determined by log-rank test (\**p*<0.05, \*\**p*<0.01).

We first analyzed tumor-infiltrating lymphocytes in the bone marrow chimera mice 14 days after tumor implantation to assess the environmental effect on T cells. We found increased CD3^+^ T cell infiltration in the female-to-female group (F to F; female control) compared to the male-to-male group (M to M; male control) in both CD8^+^ T cells and CD4^+^ T cells, but not in Foxp3^+^ cells (**Fig. 4B**), confirming that T cells in the bone marrow chimera recapitulate their behavior observed in the B6 wild-type mouse model (**Fig. 2A**). Interestingly, the frequency of infiltrating T cells in the male-to-female (M to F) group was significantly lower than in the F to F group but comparable to the M to M group, suggesting male leukocyte-intrinsic regulation. Meanwhile, the female-to-male (F to M) group showed a pattern more similar to the M to M group rather than the F to F group (**Fig. 4B)**, indicating an influence of male environment on female cells. While there was no difference in inhibitory receptor expression (**Fig. 4C)**, we found that the PEX subset was largely increased in the M to M group, with a significantly decreased EFF subset (**Fig. 4D**). No difference was observed in the TEX subset. Importantly, neither the M to F nor the F to M group showed an increase in the PEX subset, suggesting critical roles for a combined hematopoietic cell-intrinsic and cell-extrinsic effect on the higher frequency of PEX in male mice shown in **Fig. 2D**.

Next we performed survival analysis on the bone marrow chimera mice after implantation of SB28 cells to assess immune cell-intrinsic and cell-extrinsic effects on tumor growth. The control groups, M to M and F to F, replicated the survival differences (**Fig. 4E**) as observed in intact B6 mice (**Fig. 1A**). Reconstitution of female recipients with male immune cells (M to F) significantly shortened the survival of female mice similar to the M to M group, as we have previously shown (20), and the F to M group showed extended survival comparable to the F to F group. These data support that the sex of immune cell origin is a critical factor that determines tumor progression. Collectively, these findings suggest that the sex difference in anti-tumor immunity is predominantly regulated in a hematopoietic immune cell-intrinsic manner but is also subject to environmental influences.

### T cell-intrinsic regulation of sex differences

Our findings in our *in vivo* models suggested that male and female T cells are activated to develop to different functional stages during tumor progression (**Fig. 2**). Thus, we hypothesized that male and female T cells undergo distinct types of cellular reprogramming in the highly suppressive tumor microenvironment. To test this hypothesis, we induced exhaustion of T cells *in vitro* by repeated stimulation (**Fig. 5A**). Compared to the cells stimulated only once, both male and female T cells showed increased expression of the exhaustion markers PD1, TIM3, and TOX upon repeated stimulation (**Fig. 5B**, upper panel). However, intracellular cytokine levels measured by flow cytometry after polyclonal stimulation (PMA/ionomycin) showed that female exhausted T cells retained their functionality, with higher expression of IFN-γ, TNF, and granzyme B (**Fig. 5B**, lower panel). qPCR analysis confirmed that female exhausted T cells exhibited higher expression of genes encoding anti-tumor effector cytokines (*ifng, gzmb, il2)* and transcription factors related to effector functions (*tbx21, eomes*) **(Fig. 5C)**. Interestingly, transcript levels of markers associated with exhaustion status were increased in female cells (*pdcd1, havcr2*) or comparable (*tox, batf, irf4, tigit*) between male and female cells. These data suggest that male and female T cells may undergo distinct cell-intrinsic regulation of their functional state during exhaustion.

**Figure 5.**
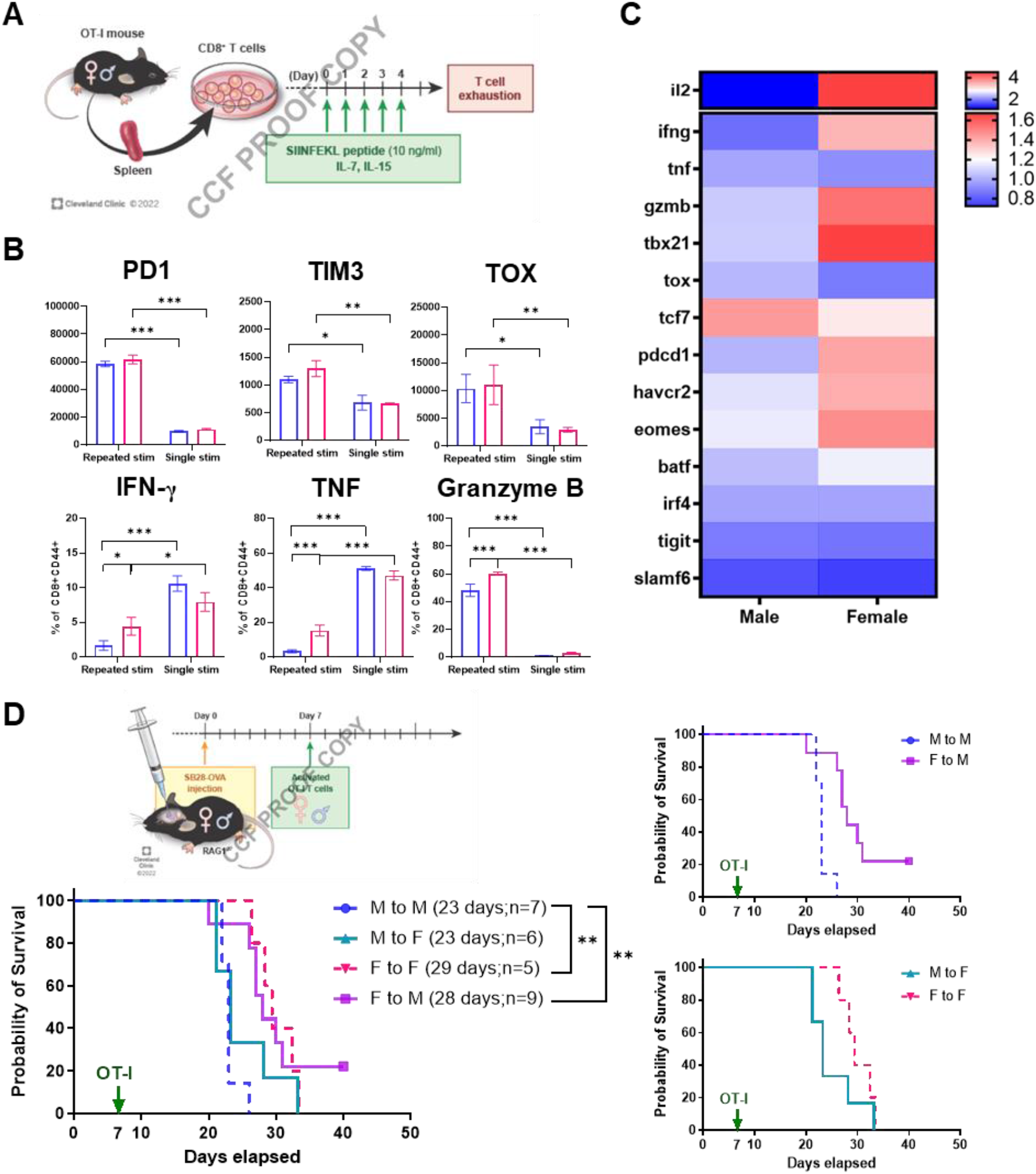
Cell-intrinsic regulation of sex differences in T cell function. (A) Schematics depicting *in vitro* generation of exhausted T cells. (B) Exhaustion markers and cytokine expression were measured on day 5 by flow cytometry after polyclonal stimulation with stimulation cocktail for 4 hours. Data shown as mean±SD and is representative of three independent experiments. Two-way ANOVA analysis with Tukey’s multiple comparison test (\**p*<0.05, \*\**p*<0.01). (C) qPCR analysis on exhausted T cells. Relative expression levels normalized to male T cells are shown. (D) Kaplan-Meier curves depicting survival of male and female RAG1^-/-^ mice bearing SB28-OVA tumor cells after adoptive transfer of OT-I cells. Data shown is combined from two independent experiments. Statistical significance was determined by log-rank test (\**p*<0.05, \*\**p*<0.01).

Next, we evaluated the intrinsic role of T cells in driving the observed sex differences in GBM survival. To avoid female T cell-mediated rejection of males, we utilized an adoptive transfer model using the OVA-OT-I system in RAG1^-/-^ mice (**Fig. 5D**; as in **Fig. 1D**). Transfer of female T cells delivered a survival advantage to recipients, as female OT-I cells significantly extended survival of male recipients compared to the male-to-male group. In contrast, male OT-I cells shortened the survival of female recipients (**Fig. 5D**). These results indicate that the ability of T cells to control tumor growth is dominantly determined by the sex of the originating host, not the recipient’s environment.

### Male-biased T cell exhaustion in GBM patients and human T cells

To investigate whether sex differences in T cells are recapitulated in human GBM patients, we analyzed exhausted CD8^+^ T cell subsets from GBM patient tumors using flow cytometry. KLRG1 and PD1 expression was used to exclude the short-lived effector T cell population, and exhausted T cells subsets were determined based on expression of TCF1, TIM3, and CXCR5 (23) (**Fig. 6A**). There was no significant difference in the percentage of CD8^+^ T cells and the PD1^+^KLRG1^-^ population from male and female tumors (**Supplementary Fig. 8A**). Meanwhile, an increased frequency of progenitor exhausted T cells (CD8^+^KLRG1^-^PD1^+^CXCR5^+^TCF1^+^TIM3^-^) was found in male compared to female tumor samples (**Fig. 6B**), while no difference was observed in the subsets of terminally exhausted T cells (CD8^+^KLRG1^-^PD1^+^CXCR5^-^TCF1^-^TIM3^+^) (**Supplementary Fig. 8B**). Additionally, increased expression of the exhaustion marker TOX was found in CD8^+^ T cells from male tumors (**Fig. 6C**), while the expression levels of other marker were comparable (**Supplementary Fig. 8C**). These results indicate that T cells from GBM patients exhibit a male-biased exhaustion status in line with our mouse model.

To further address sex differences in T cells under exhaustion conditions, we performed an *in vitro* exhaustion assay by repeatedly stimulating human CD8^+^ T cells isolated from PBMCs of healthy volunteers (**Fig. 6D**). Both male and female T cells exhibited elevated expression levels of exhaustion markers (PD1, TIM3, TOX, CD39) after repeated stimulation for 12 days compared to their singly stimulated counterparts (**Fig. 6E**). Intracellular cytokine expression analysis revealed that the ability of T cells to produce dual cytokines dramatically decreased after the second stimulation (day 6), while female CD8^+^ T cells consistently expressed higher levels of IFN-γ and TNF compared to male cells (**Fig. 6F**). Interestingly, qPCR analysis showed significantly higher expression of a set of transcription factors related to T cell exhaustion (*IRF4, TOX, TCF1, EOMES, MYC)* in female T cells on day 6, but not on day 12, which suggests differential transcriptional regulation in male and female T cells (**Supplementary Fig. 8D**).

Taken together, our findings indicate that male T cells are more prone to exhaustion, which leads to accelerated tumor growth in males but potentially provides them with a larger benefit from anti-PD1 mAb therapy, whereas female T cells tend to maintain higher functionality and protect the host from tumor progression (**Fig. 6G**).

**Figure 6.**
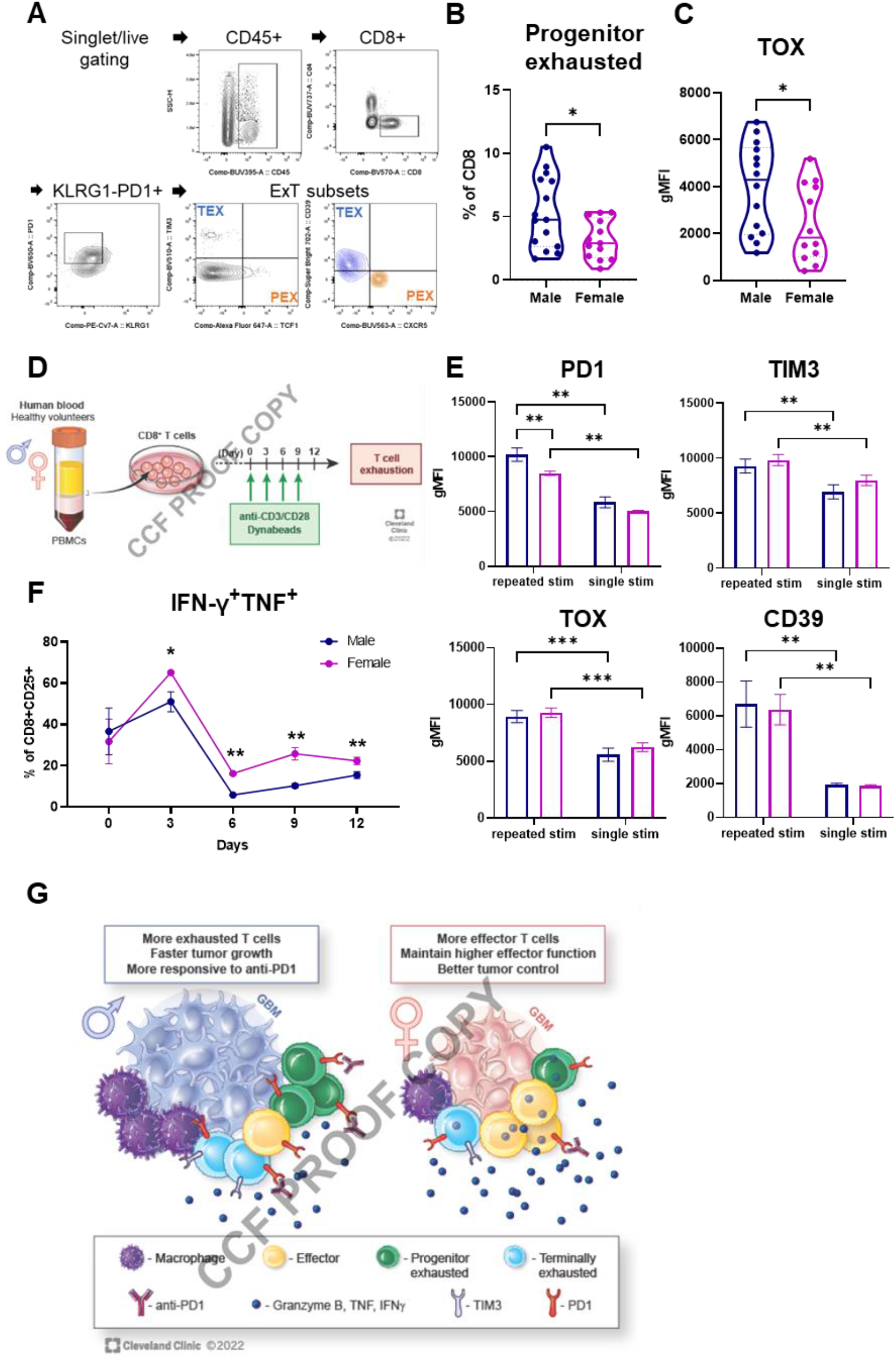
Sex differences in exhausted T cells in GBM patients. (A) A gating strategy for exhausted T cell subsets from GBM patient tumors. (B) Frequency of progenitor exhausted T cells (PEX; CD8^+^KLRG1-PD1^+^CXCR5-TCF1^+^TIM3-) and (C) TOX expression in CD8^+^ T cells from tumors of male (n=18) and female (n=14) patients with IDH-wild type GBM tumors. Unpaired t-test (\**p*<0.05). (D) *In vitro* induction of exhaustion in human CD8^+^ T cells. (E) Exhaustion marker expression in CD8^+^ T cells on day 12 post-stimulation. Two-way ANOVA analysis with Tukey’s multiple comparison test (\**p*<0.05, \*\**p*<0.01, \*\*\**p*<0.001). (F) Intracellular expression of IFN-γ^+^TNF^+^ in CD8^+^ T cells during repeated stimulation. Multiple unpaired *t*-test was performed (\**p*<0.05, \*\**p*<0.01). Data shown as mean±SD and is representative of two independent experiments. (G) Proposed model of sex-specific T cell phenotype and functionality in GBM patients.

## Discussion

Sex differences are emerging as a major contributor to cancer progression and therapeutic response through distinct genetic, epigenetic, and immunological mechanisms (24, 25). We previous reported a sex difference in MDSC localization in GBM whereby males had increased m-MDSCs in the TME while females had increased g-MDSCs in the periphery. This difference was leveraged for the development of sex-specific therapies that were validated in pre-clinical models (20). Importantly, while functional and targetable sex-differences present within certain myeloid populations, we demonstrate here that GBM-infiltrating T cells actually mediate sex difference in overall survival. Using pre-clinical models and human patient validation, we demonstrate increased T cell exhaustion in males compared to females, with males displaying an enhanced response to single-agent ICI treatment. Mechanistically, the T cell-dependent survival difference was predominantly due to hematopoietic immune cell-intrinsic differences along with the impact of environment including sex hormones.

Our observation that males are more responsive to single-agent ICI treatment, based on sex differences in T cell exhaustion and inhibitor receptor expression, supports clinical trial data in many cancers whereby males show an enhanced therapeutic response and females develop more adverse events (26-28). The latter observation is likely due to enhanced immune activation status in women. Interestingly, in lung cancer patients, females showed an enhanced response to PD1/PD-L1 when combined with chemotherapy compared to males (29), suggesting that additional stimuli are required to activate an anti-tumor immune response in females. For GBM, this will likely be needed, as ICI monotherapies have not shown strong clinical benefit and current immunotherapy strategies are now focused on combination therapies (30). Furthermore, immediate priorities currently include sex-specific assessment of combination strategies in pre-clinical models and using sex as stratification criteria for early-stage clinical trials.

Our pre-clinical data also demonstrate that male T cells are more exhausted than female T cells, which has also recently been seen in several malignancies including melanoma, colon cancer, and bladder cancer models (17, 18). These observations were also validated in human GBM patients and are consistent with reports in melanoma patients (31) but opposite to what has been observed in lung cancer patients (32). While both of these tumors are more responsive to ICIs than GBM (33, 34) and have an increased mutational burden compared to GBM (35), it is unclear why there are differences and why GBM appears to more closely phenocopy melanoma. This is likely to be due to a combination of differences in driver mutations, standard-of-care therapies, and anatomical locations. For GBM in particular, the specialized neural-immune microenvironment is likely to provide unique stimuli (36), and sex differences in other neural cell types may also impact T cell function. Future studies focusing on the interaction between brain-specific mechanisms and T cells are likely to help clarify the molecular underpinnings of sex differences in T cell function and may reveal sex-specific mechanisms that could be leveraged for next-generation therapies.

Androgen in particular has recently received attention for its role in regulating T cell exhaustion, with contradictory molecular mechanisms (17, 18). These groups have shown that blockade of androgen receptor signaling restored CD8^+^ T cell function, with increased responsiveness in anti-PD1/PD-L1 treatment in males, in line with a previous report on castration-resistant prostate cancer (37). Our *in vitro* data, however, along with our adoptive transfer studies and bone marrow chimera studies, suggest a predominant cell-intrinsic underpinning of male T cell exhaustion. We reached these conclusions based on the maintained difference in T cell exhaustion in males versus females *ex vivo*, in the absence of the influence of sex hormones. Yet this does not rule out the possibility that the initial impact of sex hormones on stem cell stage or pre-isolation have been maintained through epigenetic memory, which needs further investigation. These cell-intrinsic differences could be potentially derived from sex chromosomes via expression of genes escaping X chromosome inactivation or micro-RNAs highly enriched in X chromosomes (38). Differential expression of epigenetic regulators encoded on sex chromosomes and their roles in immune responses have been reported (39-41), suggesting the possibility of sex-specific epigenetic reprogramming of T cells. Given that sex differences in GBM are observed throughout all age groups, while sex hormone levels vary (42), delineating the effects of sex hormones and sex chromosomes in sexually dimorphic GBM immunity requires further investigation.

Sex differences in T cell responses in GBM are likely derived from a combination of sex hormone-derived influences and cell-intrinsically derived influences. The molecular drivers of these and how the two intersect should be the focus of future studies to better enable us to understand immunological sex differences and tailor therapies accordingly. Additionally, an aspect outside of the scope of this study but still of pressing interest is the extent to which sex differences in antigen-presenting cells are present and affect T cell behavior. A meta-analysis on GBM clinical trials employing autologous dendritic cells showed that female patients had a more robust survival advantage compare to male patients, providing a rationale to understand underlying mechanisms (43). Taken together, our study identifies T cells as a critical component driving sex differences in GBM progression and a male-biased T cell exhaustion state that could potentially interrogate sex-specific immunotherapy responses in cancer patients. Our study provides insight into the immunological mechanisms underlying sex differences in GBM and further emphasizes a need for sex-specific treatment strategies.

## Materials and Methods

### Cell lines

The syngeneic mouse GBM cell lines SB28 and ovalbumin-overexpressing SB28 were kindly gifted by Dr. Hideho Okada (UCSF). GL261 cells were obtained from the Developmental Therapeutic Program, National Cancer Institute (Bethesda, MD). All cell lines were maintained in complete RPMI1640 (Media Preparation Core, Cleveland Clinic) supplemented with 10% FBS (Thermo Fisher Scientific), 1% penicillin/streptomycin (Media Preparation Core) and GlutaMAX (Gibco). Cells were maintained in humidified incubators held at 37°C and 5% CO2 and not grown for more than 15 passages.

### Mice

All animal procedures were performed in accordance with the guidelines of the Cleveland Clinic Institutional Animal Care and Use Committee. C57BL/6 (RRID:IMSR_JAX:000664), B6 CD45.1 (B6.SJL-Ptprc^a^Pepc^b^/BoyJ; RRID:IMSR_JAX:002014), RAG1^-/-^ (B6.129S7-Rag1tm1Mom/J; RRID:IMSR_JAX:002216), and OT-I TCR transgenic (C57BL/6-Tg(TcraTcrb)1100Mjb/J; RRID:IMSR_JAX:003831) mice were purchased from the Jackson Laboratory as required. NSG (NOD.Cg-Prkdc^scid^Il2rg^tm1Wjl^/SzJ) mice were obtained from the Biological Research Unit (BRU) at Lerner Research Institute, Cleveland Clinic. All animals were housed in specific-pathogen-free facility of the BRU with a light-dark period of 12 h each.

For tumor implantation, 5-6 week-old mice were anesthetized, fit to the stereotaxic apparatus, and intracranially injected with 10,000-25,000 tumor cells in 5 μl RPMI-null media into the left hemisphere approximately 0.5 mm rostral and 1.8 mm lateral to the bregma with 3.5 mm depth from the scalp. In some experiment, 5 μl null-media was injected into age- and sex-matched animals for sham controls. Animals were monitored over time for the presentation of neurological and behavioral symptoms associated with the presence of a brain tumor.

In some experiments, mice were treated with anti-PD1 (BioXcell, Cat# BE0273; RRID: AB_2687796) or isotype antibody (BioXcell, Cat# BE0089, RRID:AB_1107769) intraperitoneally starting from 7 days post-tumor implantation, and injections were repeated every 2-3 days for five times.

In the adoptive transfer model, RAG1^-/-^ mice received intracranial injection of SB28-OVA tumor cells (15,000 cells/mouse). OT-I splenocytes were activated *in vitro* with 2 μg of SIINFEKL peptides (Sigma) in the presence of recombinant human IL-2 (50 U/ml, Peprotech) for 3 days. Activated CD8^+^ T cells (5×10^6^ cells/mouse) were transferred intravenously into SB28-OVA tumor-bearing mice on day 7 post-tumor implantation.

To generate bone marrow chimeras, 5-week-old B6. CD45.2 male and female mice were irradiated with 12 Gy in total given in two fractions 3-4 hours apart. Bone marrow cells were obtained from tibias and femurs of 5-week-old B6. CD45.1 mice, and existing T cells were depleted using Thy1.2 (BioXcell, Cat# BE0066, RRID:AB_1107682) and rabbit complement. A total of 5×10^6^ T cell-depleted bone marrow cells were injected retro-orbitally into the irradiated recipients. Animals were given Sulfatrim in drinking water for 2 weeks to prevent infection. After 6-8 weeks, animals were subjected to tumor implantation.

### Immunophenotyping by flow cytometry

At the indicated time points, a single-cell suspension was prepared from the tumor-bearing left hemisphere by enzymatic digestion using collagenase IV (Sigma) and DNase I (Sigma), followed by straining with a 40-μm filter. Cells were stained with LIVE/DEAD Fixable stains (Thermo Fisher) on ice for 15 min. After washing with PBS, cells were resuspended in Fc receptor blocker (Miltenyi Biotec) diluted in PBS/2% BSA and incubated on ice for 10 min. For surface staining, fluorochrome-conjugated antibodies were diluted in Brilliant buffer (BD) at 1:100 – 1:250, and cells were incubated on ice for 30 min. After washing with PBS-2% BSA buffer, cells were then fixed with FOXP3/Transcription Factor Fixation Buffer (eBioscience) overnight. For tetramer staining, cells were incubated with tetramer antibody diluted in PBS/2% BSA at 1:1000 dilution after the FcR blocker step on ice for 30 min, followed by surface staining. For intracellular staining, antibodies were diluted in FOXP3/Transcription Factor permeabilization buffer at 1:250-1:500, and cells were incubated at room temperature for 45 min. For intracellular cytokine detection, cells were stimulated using Cell Stimulation Cocktail plus protein transport inhibitor (eBioscience) in complete RPMI for 4 hours. After stimulation, cells were subjected to the staining procedures described above. Stained cells were acquired with a BD LSR Fortessa (BD) or Aurora (Cytek) and analyzed using FlowJo software (v10, BD Biosciences).

### Reagents

For immunophenotyping in mouse models, the following fluorophore-conjugated antibodies were used: CD11b (M1/70, Cat# 563553), CD11c (HL3, Cat# 612796), Ly6G (1A8, Cat# 560603), CD3 (145-2C11, Cat# 564379), CD44 (IM7, Cat# 612799) from BD biosciences. CTLA4 (UC10-4B9, Cat# 106312), PD1 (29F.1A12, Cat# 135241), B220 (RA3-6B2, Cat# 103237), Ki-67 (11Fb, Cat# 151215), TIM3 (RMT3-23, Cat# 119727), I-A/I-E (M5/114.15.2, Cat# 107606), CD45 (30-F11, Cat# 103132), LAG3 (C9B7W, Cat# 125224), NK1.1 (PK136, Cat# 108716), CD4 (GK1.5, Cat# 100422), CD8 (6206.7, Cat# 100712), Ly6C (HK1.4, Cat# 128024), CD68 (FA-11, Cat# 137024), granzyme B (QA18A28, Cat# 396413), TNF (MP6-XT22, Cat# 506329), IFN-γ (XMG1.2, Cat# 505846) were obtained from BioLegend. Anti-Foxp3 (FJK-16s, Cat# 12-5773) antibody was obtained from eBioscience. Anti-TOX (TXRX10, Cat# 12-6502) antibody was purchased from Invitrogen, and anti-TCF1 (C63D9, Cat# 6709S) antibody was obtained from Cell Signaling Technology. For the tetramer assay, anti-CD8 antibody (KT15, Cat# ab22504) was obtained from Abcam, and PE-conjugated H-2K(b) SIINFEKL tetramer was provided by the NIH Tetramer Core Facility.

For analysis of GBM patient samples, the following fluorophore-conjugated antibodies were used: CD45 (HI30, Cat# 563791), CD3 (SP34-2, Cat# 557757), CD4 (SK3, Cat# 612749), PD1 (EH12.2, Cat# 564104), CTLA4 (BN13, 561717) were obtained from BD Biosciences. CD8 (RPA-T8, Cat# 301038) and TIGIT (A15153G, Cat# 372718), KLRG1 (2F1/KLRG1, Cat# 138415), TIM3 (F38-2E2, Cat# 345030), TBET (4B10, Cat# 644817) were purchased from BioLegend. Anti-CD39 (A1, Cat# 67-0399) antibody was obtained from eBioscience. Anti-TOX and anti-TCF1 antibodies were obtained as described above.

### In vitro generation of exhausted T cells

To induce exhaustion of mouse T cells, CD8^+^ T cells were isolated from splenocytes of OT-I mice using a magnetic bead isolation kit (Stemcell Technology) and cultured following a previously published protocol (44). In brief, 10 ng of SIINFEKL peptides was added into the culture every day until day 5 (repeated stimulation), or for single stimulation, peptides were added only once on day 0, and cells were washed on day 2 and rested until day 5.

To induce exhaustion of human T cells, blood was obtained from healthy volunteers upon written consent following the Cleveland Clinic Institutional Review Board (IRB; IRB07-918). Fresh peripheral blood mononuclear cells (PBMCs) were isolated using a Ficoll density gradient, and CD8^+^ T cells were subsequently isolated using the Stemcell human CD8^+^ isolation kit (Stemcell Technology) following the manufacturer’s instructions. Induction of T cell exhaustion was performed following the previously published method with modification (45). Briefly, cells were cultured in complete RPMI containing recombinant human IL-2 (30 U/ml, Peprotech) and stimulated with anti-CD3/anti-CD28 Dynabeads (Invitrogen) at a bead:cell ratio of 1:10. Every three days, cells were harvested, washed, and cultured with fresh beads, for up to 12 days. For single-stimulated controls, cells were harvested on day 3 and cultured without further stimulation for up to 12 days.

### Real-time quantitative PCR

Total RNA was isolated using an RNeasy mini kit (Qiagen), and cDNA was synthesized using the High-capacity cDNA Reverse Transcription Kit (Applied Biosystems). qPCR reactions were performed using Fast SYBR-Green Mastermix (Thermo Fisher Scientific) on an Applied Biosystems StepOnePlus Real-Time PCR system. The threshold cycle (Ct) value for each gene was normalized to the expression levels of *gapdh* (mouse) or *ACTIN* (human), and relative expression was calculated by normalizing to the delta Ct value of one male T cell data point. Primer sequences were obtained from PrimerBank (46), and primer sequences are listed in Table S1 (mouse) and Table S2 (human).

### GBM patient samples

Cryopreserved single-cell suspension samples were obtained from the Rosa Ella Burkhardt Brain Tumor Bank in accordance with the Cleveland Clinic Institutional Review Board (IRB2559). Samples from GBM patients diagnosed as IDH (Isocitrate dehydrogenase) mutations were excluded from our study. Cells were thawed in a 37°C water bath and washed twice with warm complete RPMI. Cells were stained with LIVE/DEAD Fixable Stains for 15 min on ice and washed, followed by incubation with Fc receptor blocker (Miltenyi Biotec) for 15 min on ice. Surface marker staining was performed for 30 min on ice with following antibodies: CD45, CD3, CD4, CD8, CD44, PD1, TIM3, CD39, KLRG1, TIGIT. Cells were then fixed with FOXP3/Transcription factor fixation buffer (Invitrogen) overnight at 4°C, and intracellular staining was performed in permeabilization buffer for following markers: TBET, TCF1, CTLA4, TOX. Stained samples were acquired by Cytek Aurora and analyzed by FlowJo software.

### Statistical Analysis

GraphPad Prism (Version 9, GraphPad Software Inc. RRID:SCR_002798) software was used for data presentation and statistical analysis. Unpaired *t* test or one-way/two-way analysis of variance (ANOVA) was used with a multiple comparison test as indicated in the figure legend. Survival analysis was performed by log-rank test. *p*-value <0.05 was considered significant (\**p*<0.05, \*\**p*<0.01, \*\*\**p*<0.001).

## Author’s contributions

Conception and design: J.L., M.N., J.D.L.

Development of methodology: J.L., M.N.

Acquisition of data: J.L., D.J.S., C.L., D.B., D.C.W., A.L.

Analysis and interpretation of data: J.L., M.N., R.L.F., J.D.L.

Writing, review: J.L., M.N., D.J.S., A.D., W.G., S.J.C., A.V., R.L.F., M.S.A., J.D.L.

Administrative, technical, or material support: S.J., M.M., M.M.G., D.D.K., A.D., W.G., S.J.C., A.V., R.L.F., H.O., M.S.A., J.D.L.

Study supervision: J.D.L.

## Acknowledgements

We thank the members of the Lathia laboratory for insightful discussions. We are grateful to Drs. Jill Barnholtz-Sloan (National Cancer Institute), Josh Rubin (Washing University in St. Louis), Jim Connor (Penn State College of Medicine) and Mike Berens (TGEN) for their useful suggestions. We greatly appreciate the editorial assistance of Dr. Erin Mulkearns-Hubert (Cleveland Clinic) and illustrative work of Ms. Amanda Mendelsohn from the Center for Medical Art and Photography at the Cleveland Clinic for illustrative work. We would like to acknowledge the technical help from Cleveland Clinic Flow Cytometry Core. We also thank the NIH Tetramer Core Facility (contract number 75N93020D00005) for providing H-2K SIINFEKL tetramers. This work is supported by National Institutes of Health grants R35 NS127083 (J.D.L), P01 CA245705 (J.D.L.), K99 CA248611 (D.B.), R35NS105068 (H.O.), F30CA250254 (A.L.), T32GM007250 (A.L.), R01 HL158801 (S.J.C.), T32 5T32AI007024-40 (D.C.W), TL1 5TL1TR002549-03 (D.C.W.). This work was also supported the American Brain Tumor Association (J.D.L.), Case Comprehensive Cancer Center (J.D.L.), Cleveland Clinic/Lerner Research Institute (J.D. L.) and the American Society of Transplantation Research Network (M.N.).

