## Supplementary data for "Sex-specific T cell exhaustion drives differential immune responses in glioblastoma"

**Supplementary Materials**

Supplementary figures


Fig. S1. Mouse syngeneic GBM cell lines do not contain a Y chromosome

Fig. S2. Frequencies of tumor-infiltrating immune cells in the SB28 model

Fig. S3. No sex difference in T cells was observed at an earlier time point

Fig. S4. Male T cells are more exhausted in the GL261 model

Fig. S5. Phenotype of T cells in the periphery does not replicate the sex differences shown in tumor-infiltrating T cells

Fig. S6. PD1 blockade enhanced immune responses

Fig. S7. No significant difference in the immune cell composition of bone marrow chimera mice before tumor implantation

Fig. S8. Phenotyping tumor-infiltrating $CD8^+$ T cell from GBM patient tumors


Supplementary tables


Table S1. qPCR primers used for mouse study

Table S2. qPCR primers used for human study

**Supplementary Figure 1.**

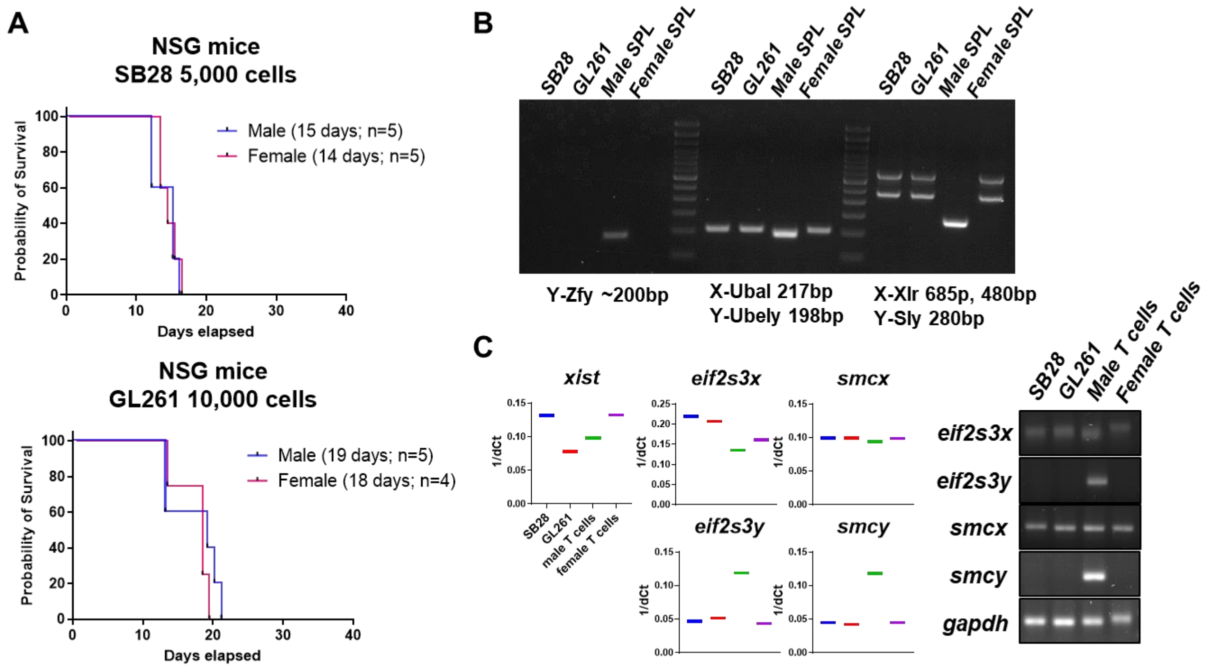


**Supplementary Figure 1. Mouse syngeneic GBM cell lines do not contain a Y chromosome.** (A) Kaplan-Meier curves depicting survival of NSG mice after intracranial injection of tumor cells (SB28, 5,000 cells; GL261, 10,000 cells). (B) PCR was performed to measure sex-specific genes (*Zfy, Ubal, Ubely, Xlr, Sly*) on genomic DNA extracted from SB28 and GL261 cells. Genomic DNA from splenocytes (SPL) of male or female mice was used as controls. Ladder size, 100 bp. (C) Expression levels of X or Y chromosome-encoded genes (*xist, eif2s3x/y, smcx/y*) were measured using qPCR. Data was normalized to GAPDH (deltaCt;dCt) and are presented as relative expression (1/dCt; left panel). Amplified transcripts were visualized on 2% agarose gel (right panel).

**Supplementary Figure 2.**

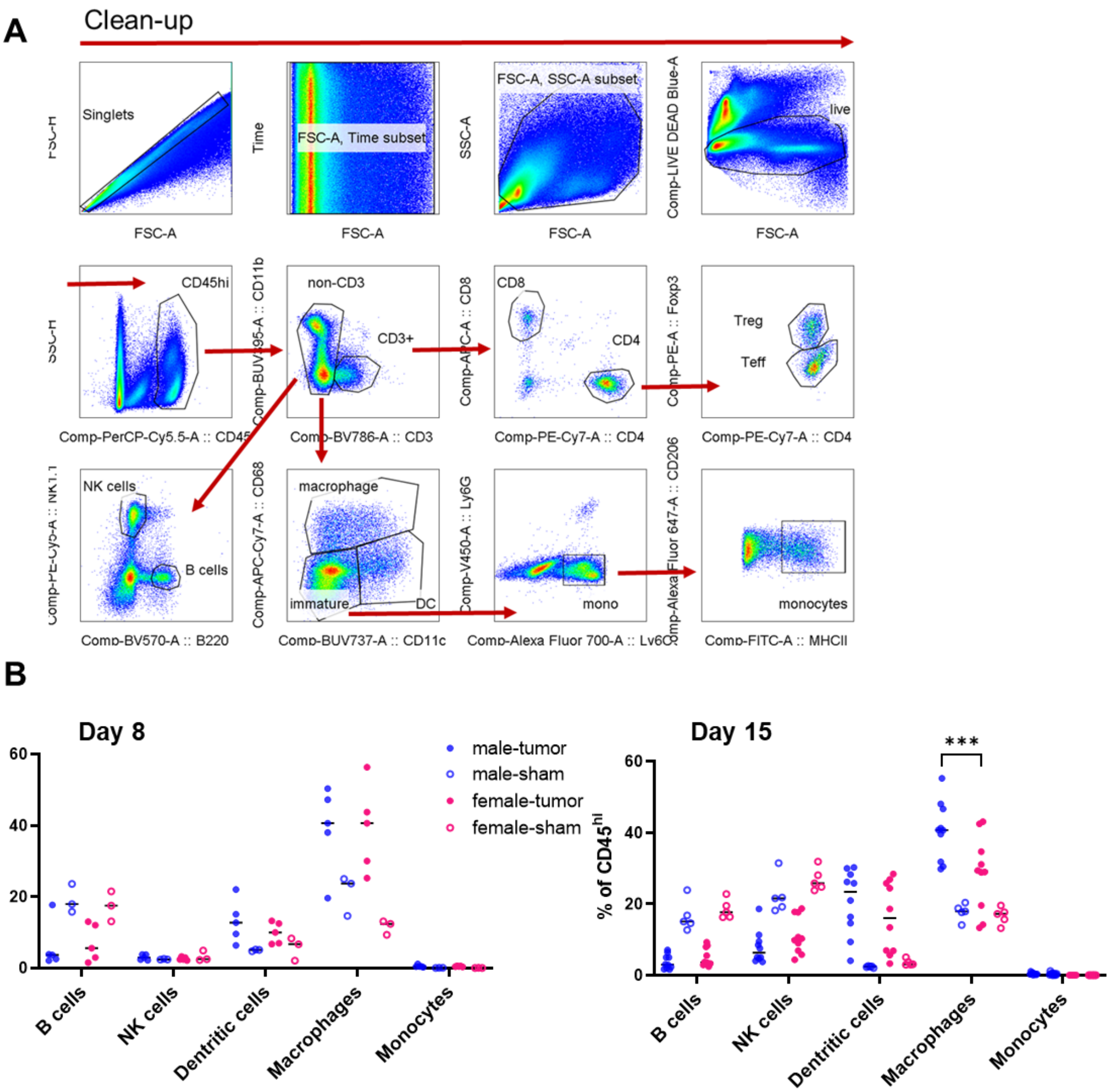


**Supplementary Figure 2**. **Frequencies of tumor-infiltrating immune cells in the SB28 model.** (A) A gating strategy of immune populations from tumors. (B) Frequency of immune cell subsets from tumors on day 8 and 15 after SB28 tumor implantation. Data shown as median of n=3-10/group. *** *p* <0.001 as determined by two-way ANOVA analysis with Tukey's multiple comparison test.

**Supplementary Figure 3.**

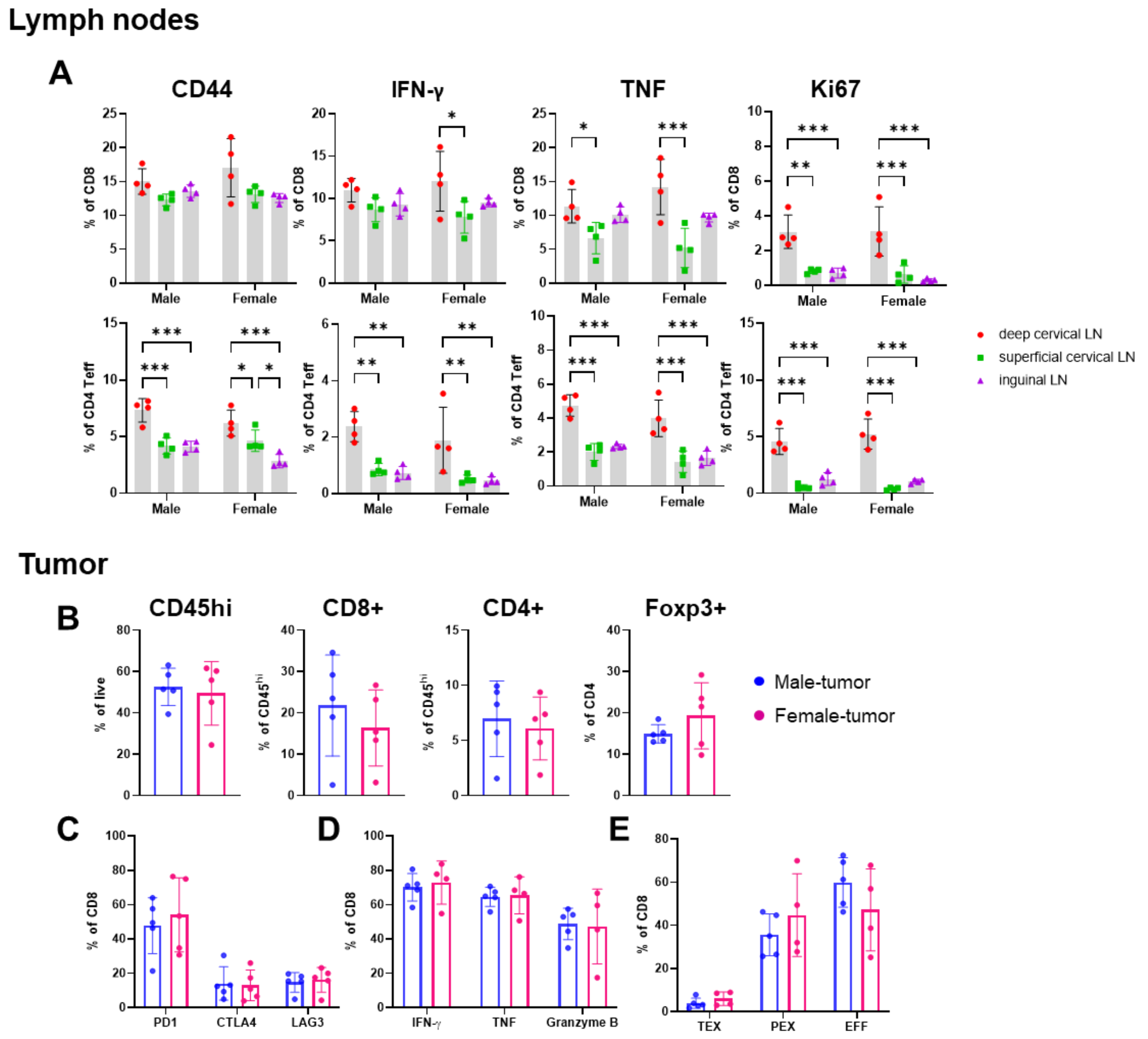


**Supplementary Figure 3. No sex difference in T cells was observed at an earlier time point.** Immune cells were analyzed on day 8 post-tumor implantation with SB28 tumor cells (15,000 cells/mouse). Data shown as mean ± SD of n=4-5/group. (A) T cell profiling on draining (deep cervical) and non-draining (superficial cervical/inguinal) lymph nodes. *$p<0.05$, ** $p<0.01$, *** $p<0.001$ as determined by two-way ANOVA with Tukey's multiple comparison test. (B) Frequency of $CD45^{hi}$ immune cells and T cell subsets in tumor. (C) Inhibitory receptor expression on $CD8^{+}$ T cells from tumors. (D) Cytokine expression was measured on $CD8^{+}$ T cells from tumors after *ex vivo* stimulation with PMA/ionomycin for 4 hours. (E) Frequency of exhausted T cell subsets and effector cells. Statistical significance was determined by unpaired *t*-test (B) or one-way ANOVA with Tukey's multiple comparison test.

**Supplementary Figure 4.**

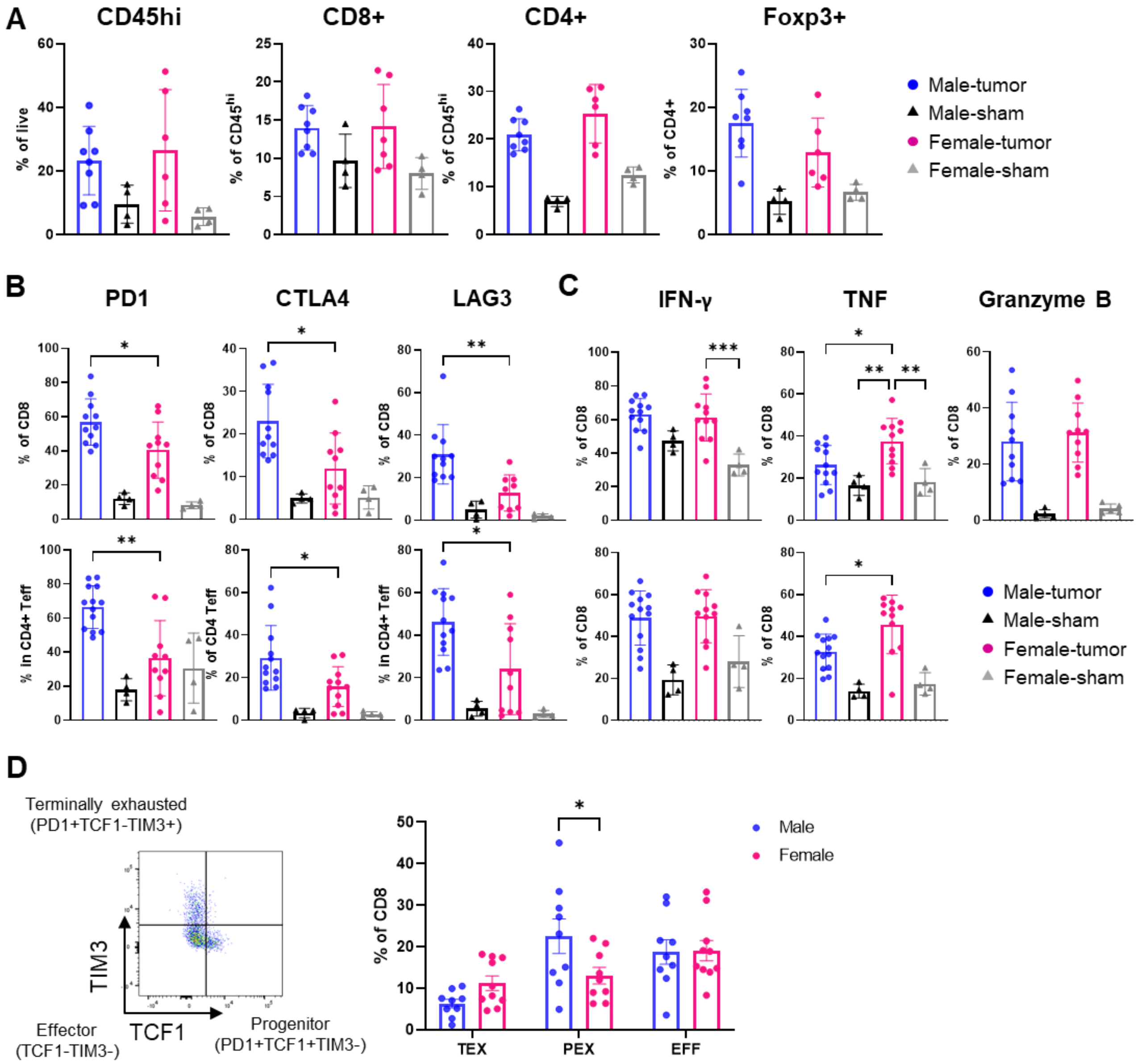


**Supplementary Figure 4. Male T cells are more exhausted in the GL261 model.** Male and female mice received intracranial implantation of 25,000 GL261 cells, and tumor-infiltrating immune cells were analyzed on day 21 post-tumor injection. Data shown as mean ± SD of n=4-12/group. (A) Frequency of CD45$^{hi}$ immune cells and T cells subsets. (B) Inhibitory receptor expression and (C) Intracellular cytokine levels were measured after ex vivo stimulation with PMA/ionomycin for 4 hours in T cell subsets. **p*<0.05, ** *p*<0.01, *** *p*<0.01 as determined by one-way ANOVA with Tukey's multiple comparison test. (D) Frequency of exhausted T cell and effector cell subsets in CD8$^{+}$ T cells. **p*<0.05, as determined by two-way ANOVA with Tukey's multiple comparison test.

**Supplementary Figure 5.**

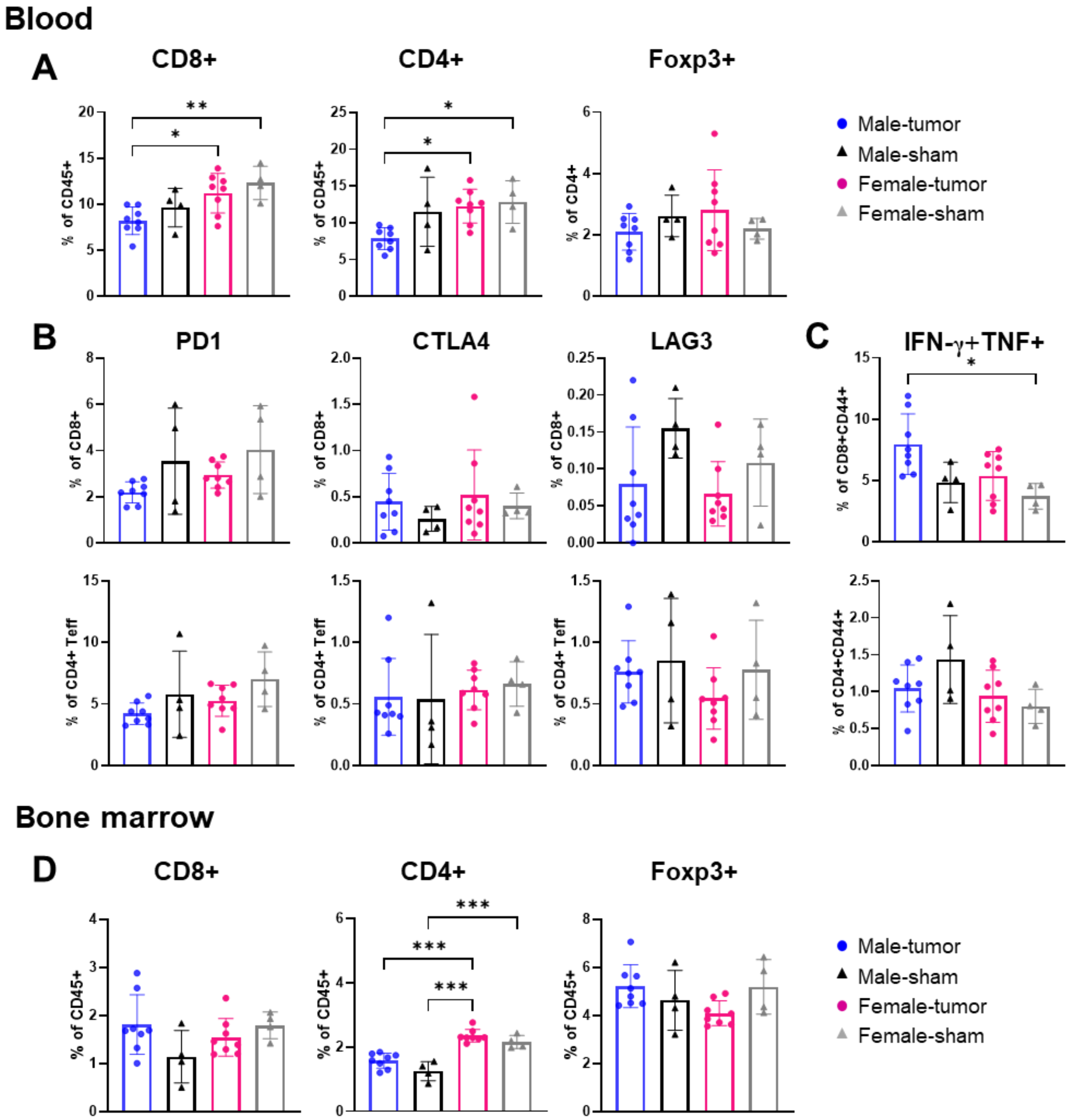


**Supplementary Figure 5. Phenotype of T cells in the periphery does not replicate the sex differences shown in tumor-infiltrating T cells.** Blood and bone marrow were collected from mice bearing SB28 tumors on day 14. Data shown as mean ± SD of n=4-8/group. **p*<0.05, ** *p*<0.01 as determined by one-way ANOVA with Tukey's multiple comparison test. (A) Frequency of T cell subsets in the blood. (B) Inhibitory receptor expression in T cell subsets. (C) Cytokine expression was measured on $CD8^+$ and $CD4^+$ T cells from blood after *ex vivo* stimulation. (D) Frequency of T cell subsets in the bone marrow.

**Supplementary Figure 6.**

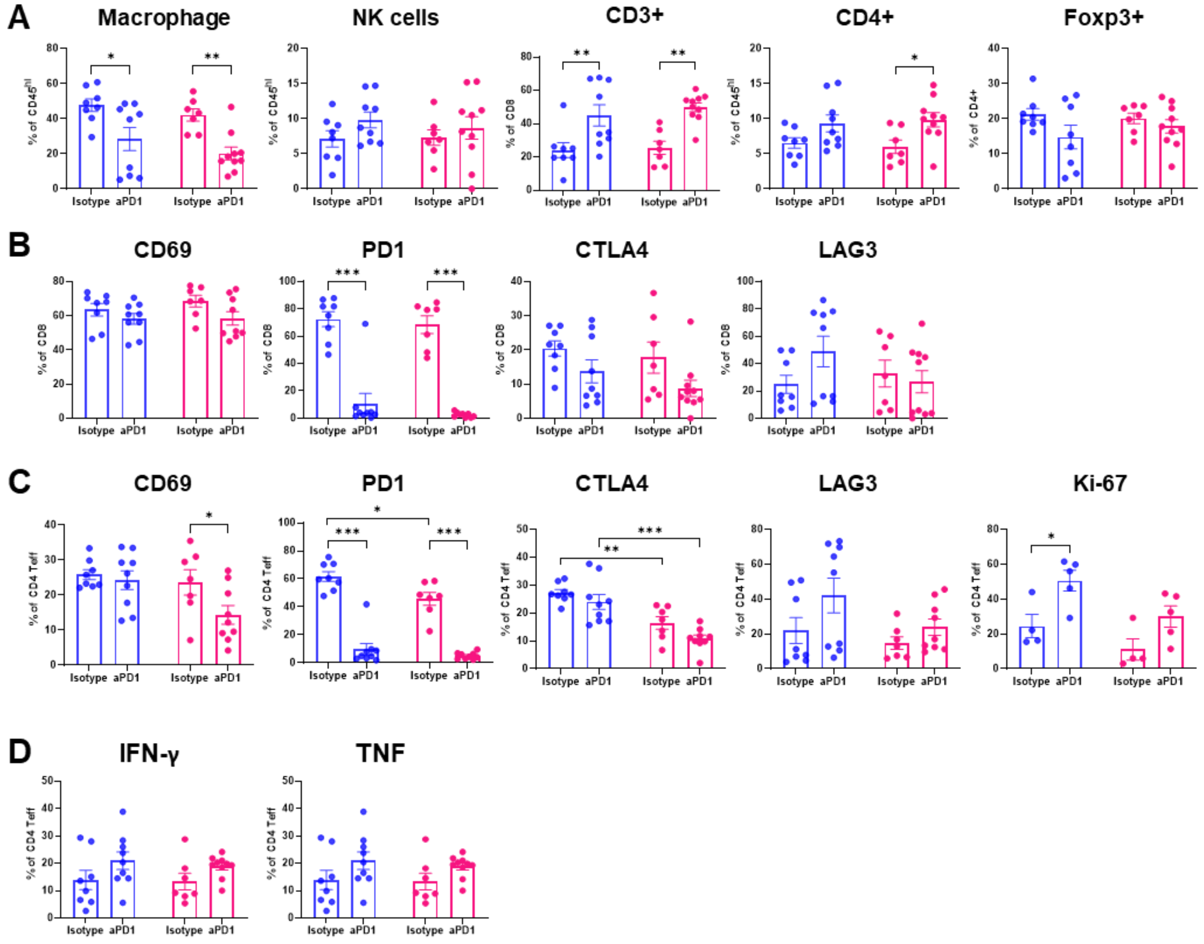


**Supplementary figure 6**. **PD1 blockade enhanced immune responses.** Mice were treated with isotype or anti-PD1 antibodies, and the tumor infiltrating immune cells were analyzed on day 18 post-tumor implantation. Data shown as mean ± SD of n=8-9/group. *$p$<0.05, ** $p$<0.01, *** $p$<0.01 as determined by two-way ANOVA with Tukey's multiple comparison test. (A) Frequency of immune cell populations. (B) Inhibitory receptor expression in CD8$^{+}$ T cells. (C) Inhibitory receptor and proliferation marker (Ki-67) expression in CD4$^{+}$Foxp3$^{-}$ T cells. (D) Intracellular cytokine expression in CD4$^{+}$Foxp3$^{-}$ T cells.

**Supplementary figure 7.**

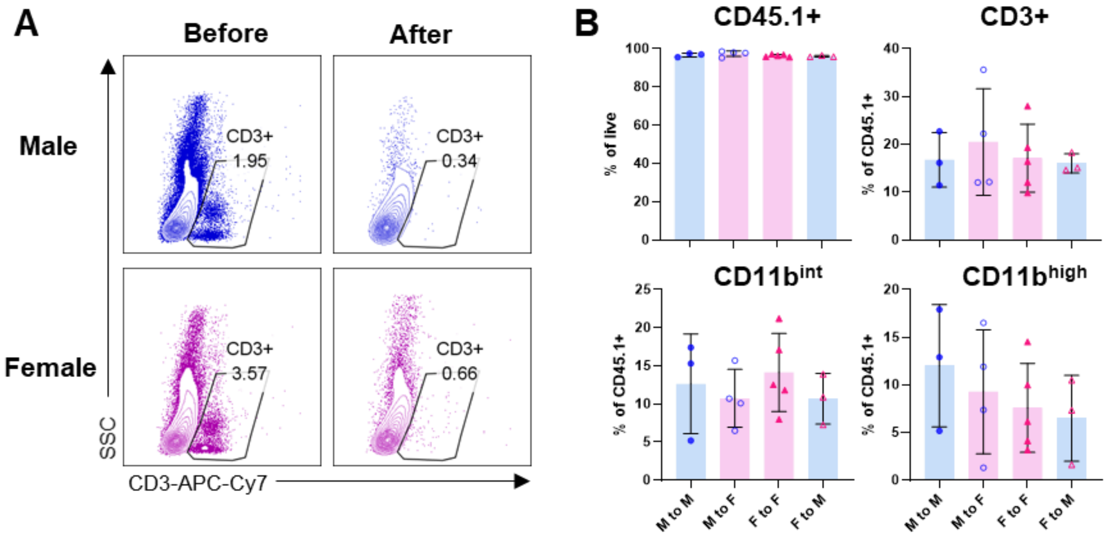


**Supplementary figure 7. No significant difference in the immune cell composition of bone marrow chimera mice before tumor implantation.** (A) Depletion of T cells before bone marrow transfer was confirmed by flow cytometry. (B) Frequency of $CD45.1^{+}$ cells and immune populations in blood was measured in bone marrow chimera mice 6 weeks after bone marrow transplantation. Data shown as mean ± SD of n=3-5/group. *$p$<0.05 as determined by one-way ANOVA with Tukey's multiple comparison test.

**Supplementary figure 8.**

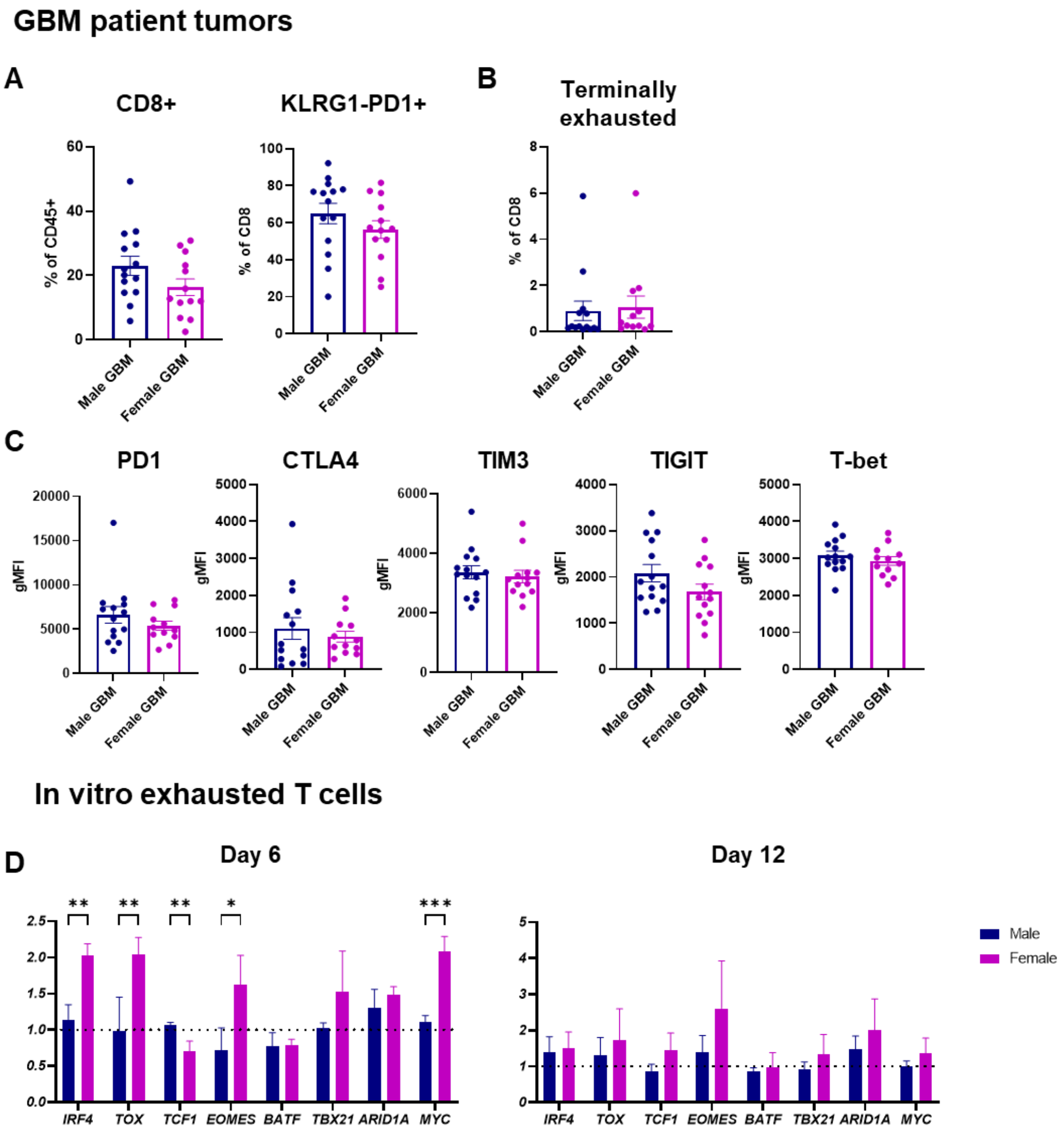


**Supplementary figure 8. Phenotyping tumor-infiltrating CD8+ T cell from GBM patient tumors**. Data is obtained from male (n=18) and female (n=14) patients with IDH-wild type GBM tumors. (A) Percentage of CD8$^+$ T cells in CD45+ immune cells and KLRG1$^-$PD1$^+$ cells in CD8$^+$ T cells. (B) Frequency of terminally exhausted T cells (TEX; CD8$^+$KLRG1$^-$PD1$^+$CXCR5$^+$TCF1$^-$TIM3$^+$) from tumors. (C) Inhibitory receptor expression in CD8$^+$ T cells. (D) mRNA expression level of transcription factors in *in vitro* exhausted T cells. Relative expression was calculated by normalization to one male sample at each time point. Multiple *t*-test was performed to determine statistical significance (**p*<0.05, ** *p*<0.01, *** *p*<0.01).

**Table S1. Primer sequences used for qPCR analysis of mouse genes.**

| Gene name | Forward primer (5’->3’) | Reverse primer (3’->5’) | Primer bank ID |
|---|---|---|---|
| *irf4* | TCCGACAGTGGTTGATCGAC | CCTCACGATTGTAGTCCTGCTT | 7305519a1 |
| *tox* | GCTCCCGTTCCATCCACAAA | TCCCAATCTCTTGCATCACAGA | 31543885a1 |
| *tcf7* | AGCTTTCTCCACTCTACGAACA | AATCCAGAGAGATCGGGGGTC | 6678245a1 |
| *eomes* | GCAATAAGATGTACGTTCACCCA | GCAGAGACTGCAACACTATCAT | 258645095c3 |
| *batf* | CTGGCAAACAGGACTCATCTG | GGGTGTCGGCTTTCTGTGTC | 7949007a1 |
| *slamf6* | ACTCCGCCTGTCAGAGGAT | AACGCCATTCATTAGCTGGGG | 133892889c1 |
| *gzmb* | CCACTCTCGACCCTACATGG | GGCCCCCAAAGTGACATTTATT | 7305123a1 |
| *havcr2* | TCAGGTCTTACCCTCAACTGTG | GGCATTCTTACCAACCTCAAACA | 160333698c1 |
| *tigit* | GAATGGAACCTGAGGAGTCTCT | AGCAATGAAGCTCTCTAGGCT | 226423923c2 |
| *pdcd1* | ACCCTGGTCATTCACTTGGG | CATTTGCTCCCTCTGACACTG | 6679239a1 |
| *tbx21* | AGCAAGGACGGCGAATGTT | GGGTGGACATATAAGCGGTTC | 9507179a1 |
| *il2* | TGAGCAGGATGGAGAATTACAGG | GTCCAAGTTCATCTTCTAGGCAC | 1504135a1 |
| *ifng* | ATGAACGCTACACACTGCATC | CCATCCTTTTGCCAGTTCCTC | 33468859a1 |
| *tnf* | GACGTGGAACTGGCAGAAGAG | TTGGTGGTTTGTGAGTGTGAG | 202093a1 |
| *gapdh* | AGGTCGGTGTGAACGGATTTG | TGTAGACCATGTAGTTGAGGTCA | 6679937a1 |

**Table S2. Primer sequences used for qPCR analysis of human genes.**

| Gene name | Forward primer (5'->3') | Reverse primer (3'->5') | Primer Bank ID |
|---|---|---|---|
| *IRF4* | GCTGATCGACCAGATCGACAG | CGGTTGTAGTCCTGCTTGC | 305410879c1 |
| *TOX* | GTGATGCCAGATATACGAAACCC | AGCTGTGACTGGTTAATGGTAGT | 51477724c3 |
| *TCF7* | CTGGCTTCTACTCCCTGACCT | ACCAGAACCTAGCATCAAGGA | 293651581c1 |
| *EOMES* | GTGCCCACGTCTACCTGTG | CCTGCCCTGTTTCGTAATGAT | 22538469c2 |
| *BATF* | TATTGCCGCCCAGAAGAGC | GCTTGATCTCCTTGCGTAGAG | 54453563a1 |
| *TBX21* | GGTTGCGGAGACATGCTGA | GTAGGCGTAGGCTCCAAGG | 7019548c1 |
| *ARID1A* | GGCGGGACTAACCCATACTC | GGCCCTGTTGACCATACCC | 117968607c3 |
| *MYC* | GGCTCCTGGCAAAAGGTCA | CTGCGTAGTTGTGCTGATGT | 239582723c1 |
| *ACTB* | CATGTACGTTGCTATCCAGGC | CTCCTTAATGTCACGCACGAT | 4501885a1 |
